# Efficient Fisher Information Computation and Policy Search in Sampled Stochastic Chemical Reaction Networks through Deep Learning

**DOI:** 10.1101/2023.04.13.535874

**Authors:** Quentin Badolle, Gabrielle Berrada, Mustafa Khammash

## Abstract

Markov jump processes constitute the central class of Chemical Reaction Network models used to account for the intrinsic stochasticity observed in the dynamics of molecular species abundance throughout Molecular Biology. These models are specified in a parametric form, and their identification requires the use of inference procedures, and in particular the estimation of the Fisher Information. Here, a fast and accurate computation method is introduced in the case of partial observations at discrete time points, based on the use of a Mixture Density Network. We also demonstrate how this Neural Network can be used to perform fast policy search. The efficiency of these approaches is illustrated on a set of examples, and is compared to that of the current state-of-the-art.

## 1 Introduction

The Stochastic Chemical Reaction Network modeling scheme is increasingly used to investigate and control cell systems in a growing number of cybergenetic platforms [1]. In these set-ups, quantifying the information content of each observation and sampling strategy is key to drive the characterisation process, and the Fisher Information is a well-established measure of it [2]. Most existing methods to compute this quantity for sampled, partially observed systems rely on the knowledge of the probability mass function and the sensitivity of the likelihood. The current state-of-the-art estimates both using the Finite State Projection [3]. The accuracy of approaches based on this approximation suffers from the degradation of the estimates with time, as well as from the curse of dimensionality when the cardinality of the truncated state-space increases [4]. In addition, the correctness of the approximated mass functions is not guaranteed to transfer to sensitivities, and no error bound has been provided in this case thus far. Finally, evaluating the Fisher Information for different sets of parameter configurations, as done in Optimal Experimental Design, implies incurring the computational cost of repeatedly solving a usually high-dimensional Differential Equation. An alternative is to first approximate the process by continuous-state dynamics, for which the Fisher Information can be computed more easily [5].

Recently developed Neural Networks offer a new way of computing the probability mass function of Chemical Reaction Networks [6, 7, 8]. As in many other fields, the introduction of Neural Networks has been motivated by their universal approximation property, their generalisation ability and their compositional structure [9]. Taking advantage of these strengths, Nessie has been shown to accurately estimate the one-dimensional marginal distributions of a range of complex, nonlinear Chemical Reaction Networks [6]. At the same time, a Deep Learning-based approach to compute the sensitivity of the likelihood and the Fisher Information is still missing.

Here, we introduce a method to compute the Fisher Information of partially observed, sampled Stochastic Reaction Networks by leveraging the compositional structure of Neural Networks. Having trained a Mixture Density Network derived from Nessie, we take its gradients with respect to its inputs, rather than its weights, and use that to estimate the sensitivities of the Chemical Reaction Network, and ultimately its Fisher Information, for arbitrary times, system parameters and even control parameters. We also demonstrate how to use the Mixture Density Network to perform policy search when the aim is to control a Stochastic Reaction Network at specific time points [10]. In this case, estimated sensitivities are used to approximate the gradient of the performance index, which is then used to perform an optimisation routine by Projected Gradient Descent.

In the rest of the paper, we first introduce our method to compute the Fisher Information of Chemical Reaction Networks using Mixture Density Networks. A set of examples has been selected to compare the performance of our approach with analytical ground truths, as well as to parallelise the initial study introducing the Finite State Projection-based method [3]. We demonstrate on these examples that the accuracy of the Deep Learning method is comparable to that of the state-of-the-art, while being faster when performed repeatedly. We then show how the sensitivities estimated from the Neural Network can also be used to perform fast policy search.

## 2 Methods

For clarity of exposition, the methods are introduced in the absence of control inputs whenever possible, and the adjustments required by the controlled case are highlighted in section S8. More generally, the details of our method are explained extensively in the Supplementary Information.

Let us consider a Stochastic Chemical Reaction Network with *N* ∈ *ℕ*^*^ molecular species, labelled *S*_*i*_ for *i* ∈ ⟦1, *N* ⟧and *M* ∈ *ℕ*^*^ chemical reactions. The jump process 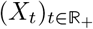 with values in ℕ^*N*^ models the abundance of species over time, and is considered over the finite horizon (0, *t*_*f*_]. Except stated otherwise, the dynamics of each reaction follow the law of mass-action.

Let us introduce the likelihood *p* : 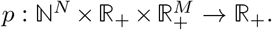 When 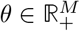 is set, 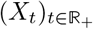 has 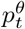 as its mass function, whose evaluation at the point *x* ∈ ℕ^*N*^ we still write *p*(*x*; *t, θ*) to emphasise its dependency on *t* and *θ*.

Whenever the state-space of 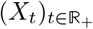 is finite, it can be projected into a subset of ℕ whose cardinality is written *N*_max_ ∈ *ℕ*^*^. We write the ordered elements of the projected state-space as 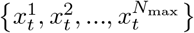. For ease of notation, we then introduce :

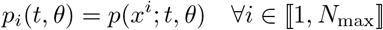

We define the sensitivity 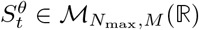 of the likelihood of *X*_*t*_ as:

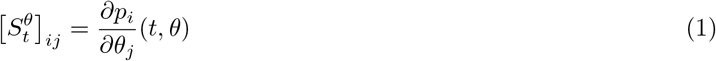

Our first aim is to compute the Fisher Information 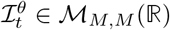 of *X*_*t*_, which we define as in classical Statistics as [3]:

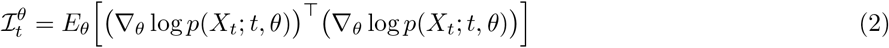

In the formula above, as well as in all other expressions involving the Fisher Information, we implicitly restrict *p* to its support.

Let us introduce a number *L* of time windows and a non-negative, piecewise-constant control input *ξ*, taking the value *ξ*_*l*_ ∈ *ℝ* over the time window *l*. Our second aim is to optimise a performance index 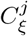 defined as:

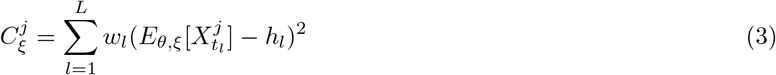

where 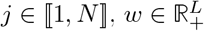 and 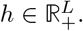.

Policy search is performed here by Projected Gradient Descent. It requires the computation of the partial derivative of 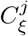 with respect to *ξ*_*i*_ for *i* ∈ ⟦1, *L* ⟧ which is given by:

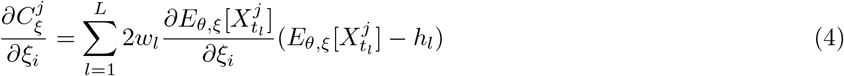

To each expectation in equations (2) and (4) corresponds an infinite sum over the state-space. Both the Deep Learning method, introduced here, and the state-of-the art, based on the Finite State Projection, approximate the mass function and the sensitivity of the likelihood as 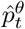 and 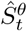 to numerically compute these sums (see sections S7 and S9). To do so, the Deep Learning method relies on a fully-connected Neural Network, which outputs mixture parameters used to define a Negative Binomial Mixture Model. The Neural Network is trained on estimated distributions of 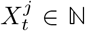 generated by the Stochastic Simulation Algorithm [11]. Each element of 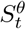 is approximated using automatic differentiation (see section S5). As for the Finite State Projection method, it defines a process with a truncated state-space for which the mass function and the sensitivity of the likelihood solve a finite-dimensional Differential Equation (see section S6).

## 3 Results

### 3.1 Computation of the Fisher Information of Chemical Reaction Networks

We first address the aim of estimating the Fisher Information of sampled, partially observed Chemical Reaction Networks. In the study introducing the Finite State Projection method for the computation of the Fisher Information, three Chemical Reaction Networks are used to demonstrate the accuracy of the approach [3]. In this subsection, we use the same examples to demonstrate that our Deep Learning-based method offers a similar performance, thereby opening a new route to fast computation of the Fisher Information and Optimal Design.

#### 3.1.1 Production and Degradation Chemical Reaction Network

We first consider the Production and Degradation Chemical Reaction Network, specified by [3]:

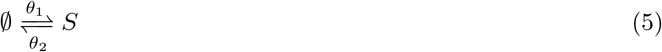

This Reaction Network has a known likelihood, from which an explicit formula for the Fisher Information can be obtained (see section S10).

The Deep Learning method introduced in this study relies on estimates of the mass function and the sensitivity of the likelihood to compute the Fisher Information. We first confirm that the Negative Binomial Mixture Model can successfully approximate the likelihood of the Production and Degradation Reaction Network by comparing it to the ground truth. The Küllback-Leibler Divergence obtained over the whole testing datasets equals 7 *·*10^−4^ on average, and the Hellinger Distance 10^−2^. This overall accuracy is further illustrated graphically by direct comparison with the ground truth in subsection S11.2.

We then investigate the accuracy of the corresponding estimated sensitivities, again using explicit formulas as references. In figure 1, sensitivities with respect to *θ*_1_ at four sampling times are displayed. All training outcomes of the Mixture Density Network are in close agreement with the reference. More specifically, they both capture the shape and values of the ground truth, as well as their overall conservation across times. Sensitivities are known to sum up to zero, and checking this offers both an indirect confirmation of the accuracy of the method, as well as a principled criterion to determine the size of the state-space to consider when subsequently computing the Fisher Information.

**Figure 1.**
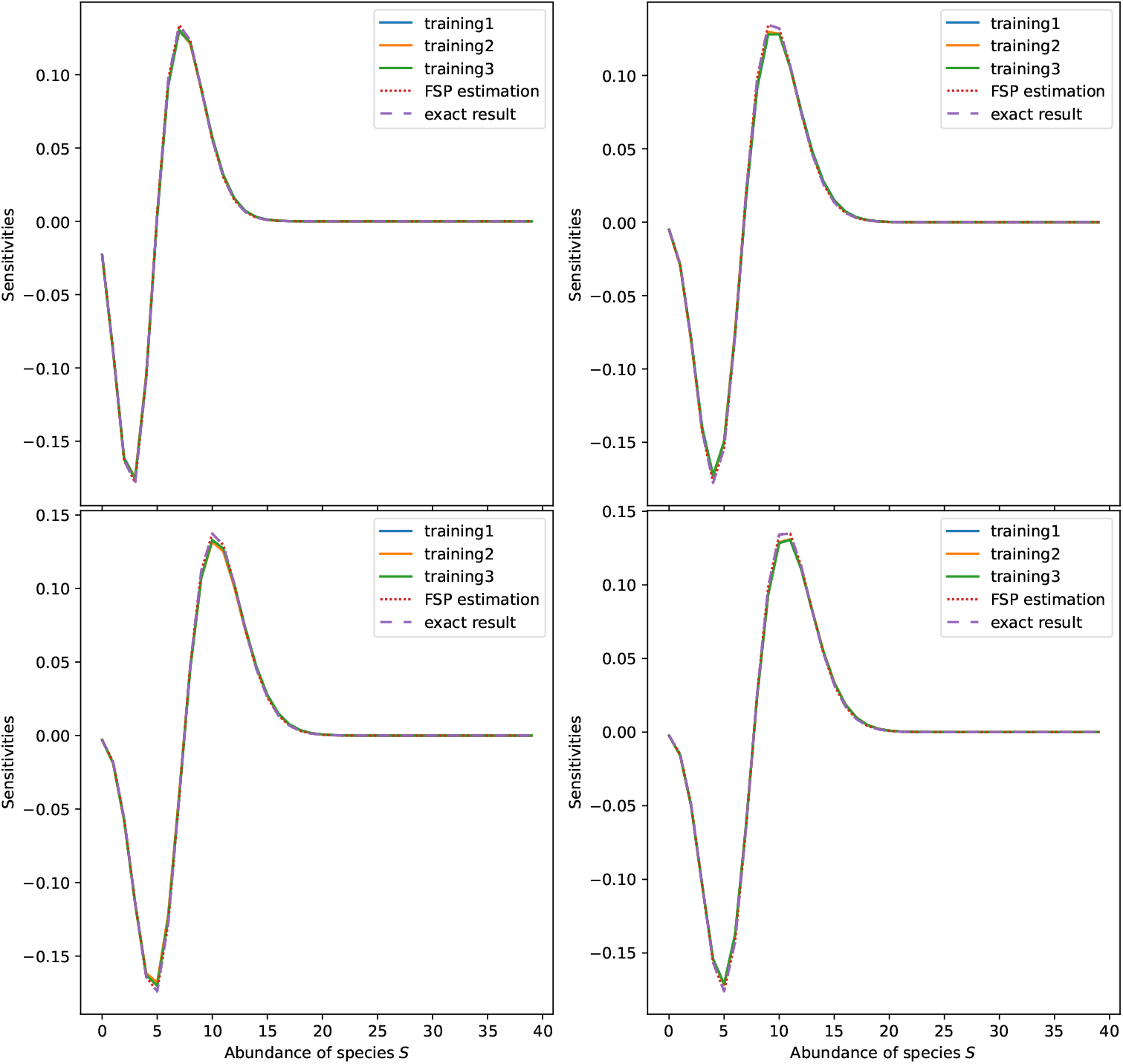
Sensitivities of the likelihood with respect to parameter *θ*_1_ for the Production and Degradation Chemical Reaction Network. Parameters: *θ*_1_ = 1.5665, *θ*_2_ = 0.1997. From left to right, top to bottom, plots correspond to times 5, 10, 15, 20.

As mentioned in the Introduction, a common motivation for using Neural Networks besides their convenient compositional structure is their generalisation property. In order to assess this ability in the present case, we explore the performance of the Mixture Density Network at a time 1.5 times greater than the maximal training time (see figure 2). We note that even at such a time outside of the training range, the Neural Network accurately predicts sensitivities.

**Figure 2.**
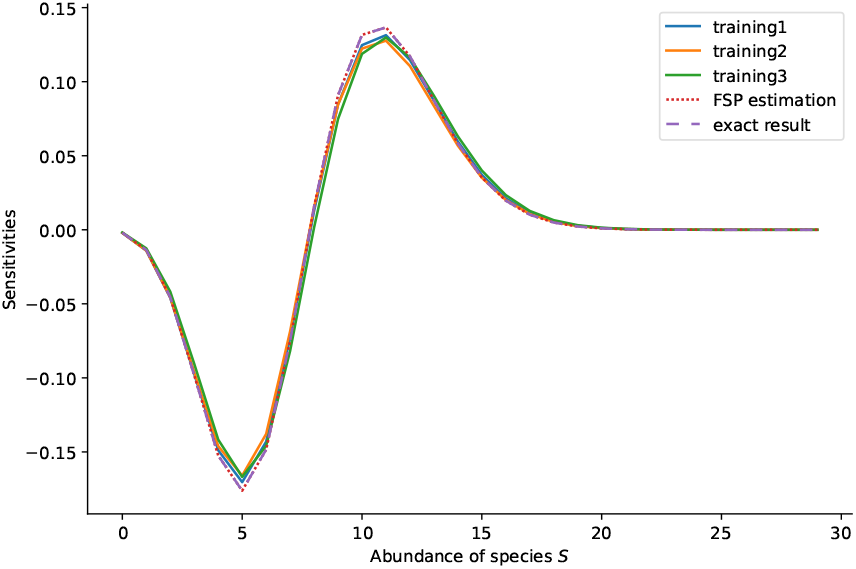
Sensitivities of the likelihood with respect to parameter *θ*_1_ for the Production and Degradation Chemical Reaction Network. Parameters: *θ*_1_ = 1.5665, *θ*_2_ = 0.1997. Time *t* = 30 is outside of the training range.

Having reliable estimates of both the likelihood and its sensitivity, we move on to the computation of the Fisher Information (figure 3). One can observe that, even for parameters never seen by the Neural Network, the relative error of the Deep Learning approach never exceeds 3.5%. Additionally, for a time markedly out of the training range, the Neural Network is able to extrapolate the knowledge learned with a relative error lower than 7%.

**Figure 3.**
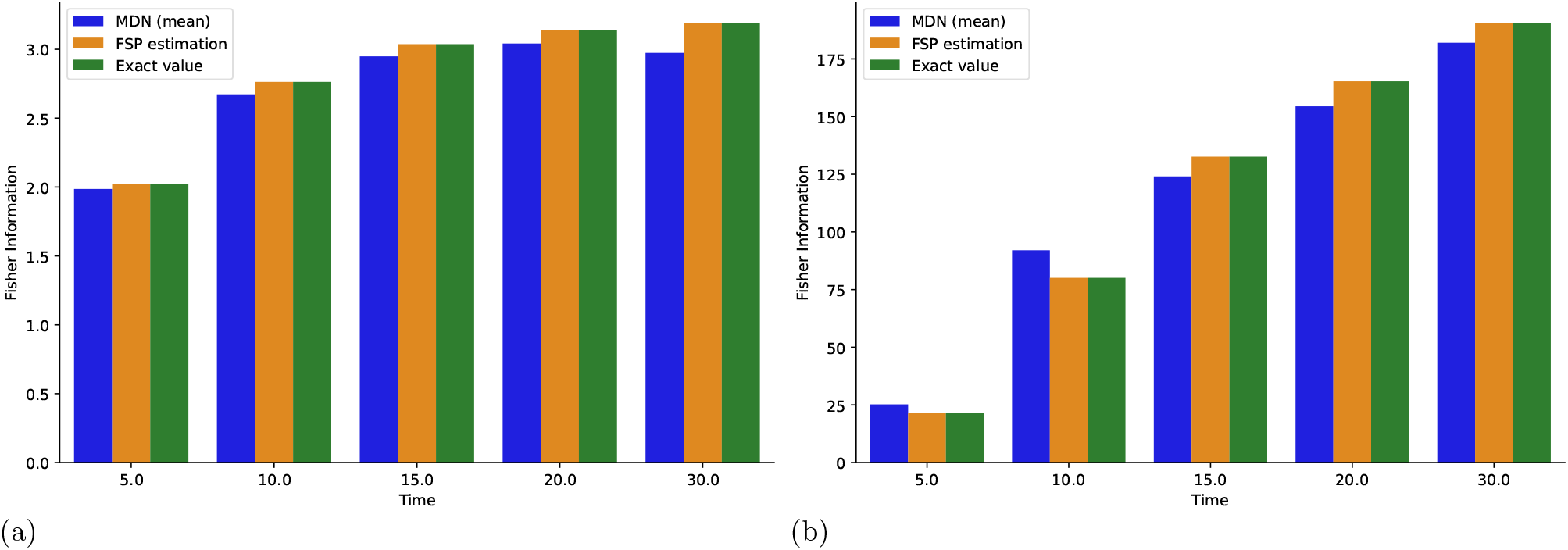
Elements 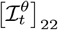 in figure 3a and 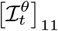 in figure 3b of the Fisher Information as a function of time for the Production and Degradation Chemical Reaction Network. Parameters: *θ*_1_ = 1.5665, *θ*_2_ = 0.1997. Time *t* = 30 is outside of the training range.

As illustrated in the second part of [3], having a method to compute the Fisher Information immediately opens the door to Optimal Experimental Design. In this context, this quantity often needs to be computed an extremely large number of times in order to identify the parameters which maximise the informational content of experiments (e.g., 69 *·*10^4^ configurations considered in [12]). Time efficiency therefore emerges as a critical performance criterion in addition to accuracy. The Deep Learning method essentially incurs a fixed, initial cost for simulation and training. Once the Neural Network has been trained, evaluation of the likelihood, its sensitivity and the Fisher Information comes at an almost marginal cost. On the other hand, for the Finite State Projection, the cost grows linearly with the number of parameter configurations *θ* and diagonal elements of the Fisher Information considered. Already when computing only one such element repeatedly, the break-even value between the two methods happens at around 7.9 *·*10^3^ parameter configurations considered ((1h06min + 5min04s)*/*0.539s, see subsection S11.1), which again is not uncommon for grid search Optimal Design.

#### 3.1.2 Bursting Gene Chemical Reaction Network

We now study the second Chemical Reaction Network used in [3]. The Bursting Gene Reaction Network is specified by:

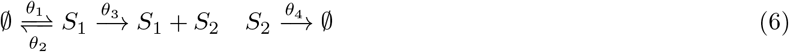

where:

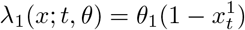

This second Reaction Network offers an occasion to explore a more challenging test bed for the Mixture Density Network, with both more species and reactions. We focus on the dynamics of *S*_2_.

This Chemical Reaction Network does not have a known likelihood, exemplifying the challenge of obtaining explicit expressions for Chemical Reaction Networks of even very moderate complexity. For the mass function, we will use distributions derived from simulations as the ground truth. As for the Fisher Information, in [3], the accuracy of the Finite State Projection method is supported by an indirect argument, which essentially states that its predictions do not refute its accuracy. We will therefore use the Finite State Projection process as a tentative point of comparison when computing the Fisher Information.

When it comes to accuracy, the general conclusions overwhelmingly follow those drawn on the previous example. The distribution predicted by the Neural Network closely matches the ground truth (see subsection S11.3). Interestingly here, the general shape of the mass function drastically changes over time. Good agreement with the reference is also observed for predicted sensitivities (figure 4) and the Fisher Information (figure 6), even for times markedly out of the training range (figures 5 and 6). Indeed, here, for the parameter *θ*_1_, the relative error in estimation at time *t* = 30 is only 0.3%.

**Figure 4.**
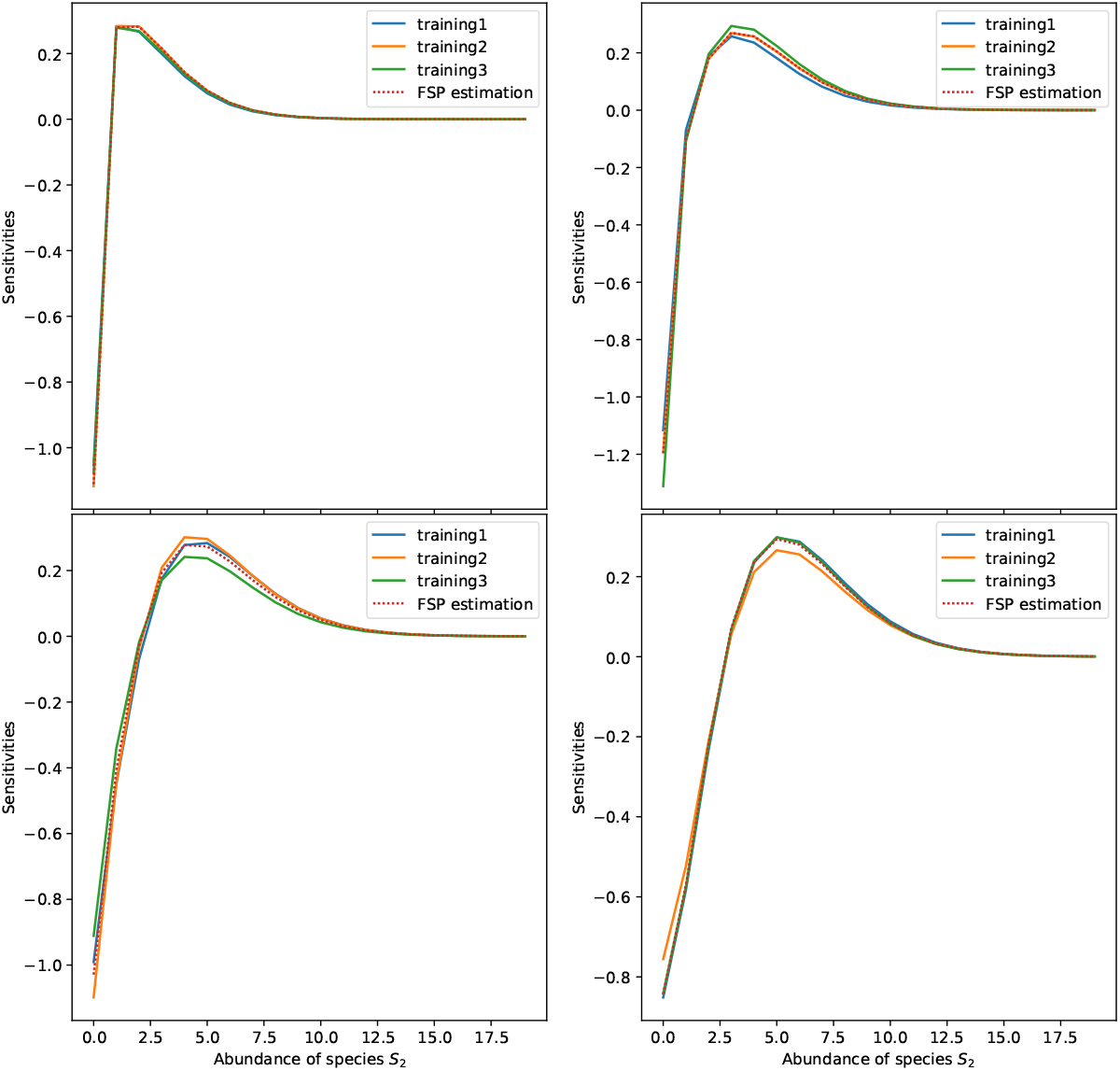
Sensitivities of the likelihood with respect to parameter *θ*_1_ for the Bursting Gene Chemical Reaction Network. Parameters: *θ*_1_ = 0.6409, *θ*_2_ = 2.0359, *θ*_3_ = 0.2688, *θ*_4_ = 0.0368. From left to right, top to bottom, plots correspond to times 5, 10, 15, 20.

**Figure 5.**
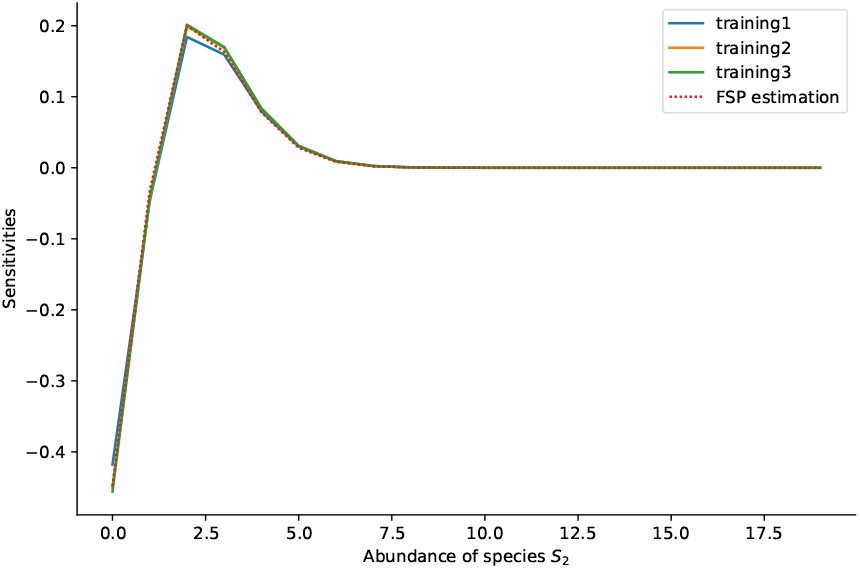
Sensitivities of the likelihood with respect to parameter *θ*_1_ for the Bursting Gene Chemical Reaction Network. Parameters: *θ*_1_ = 0.6409, *θ*_2_ = 2.0359, *θ*_3_ = 0.2688, *θ*_4_ = 0.0368. Time *t* = 30 is outside of the training range.

**Figure 6.**
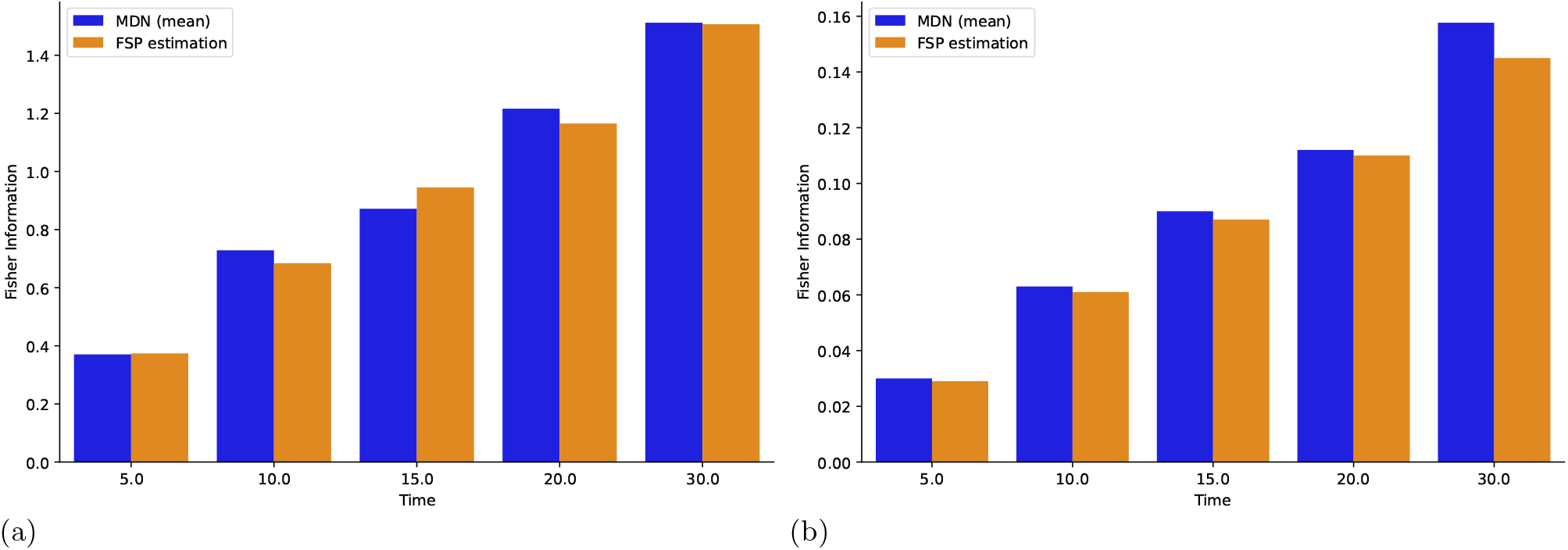
Elements 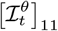 in figure 6a and 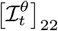 in figure 6b of the Fisher Information as a function of time for the Bursting Gene Chemical Reaction Network. Parameters: *θ*_1_ = 0.6409, *θ*_2_ = 2.0359, *θ*_3_ = 0.2688, *θ*_4_ = 0.0368. Time *t* = 30 is outside of the training range.

We are now in a position to address the topic of of computational efficiency a second time. Although the Bursting Gene Chemical Reaction Network only has one more species and two more reactions, the pivotal value at which the Deep Learning method outcompetes the Finite State Projection approach is now only 83 parameter configurations (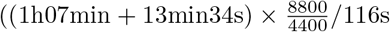, see subsection S11.1), a ∼ 95-fold reduction over the figure given in the first example.

#### 3.1.3 Toggle Switch Chemical Reaction Network

We finally investigate the last Chemical Reaction Network considered in [3], the Toggle Switch specified by:

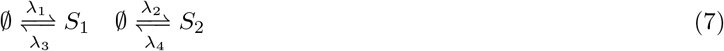

where:

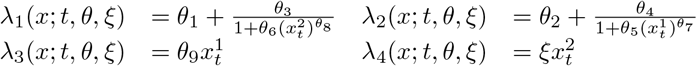

The Toggle Switch Reaction Network adds an additional layer of complexity, with non-linear propensities dependent on more than one parameter, and a reaction controlled by a time-varying parameter *ξ*.

We focus again on the dynamics of *S*_2_, and use simulations and the Finite State Projection method as ground truths for the estimation of the mass function and the sensitivities.

Similarly as before, the sensitivities of the likelihood with respect to the constant parameters agree well with those estimated by the Finite State Projection process (see figures 7a and 7b). As for the Fisher Information, the relative error of the values for parameters *θ*_2_ and *θ*_4_ never exceeds 12.8% (see figures 7c and 7d). In terms of time efficiency, the pivotal value is now attained after the computation of 1194 parameter configurations 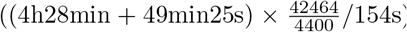. This moderate increase with respect to the previous uncontrolled Chemical Reaction Network is due to a larger training dataset required to reflect the impact of the time-varying parameters.

**Figure 7.**
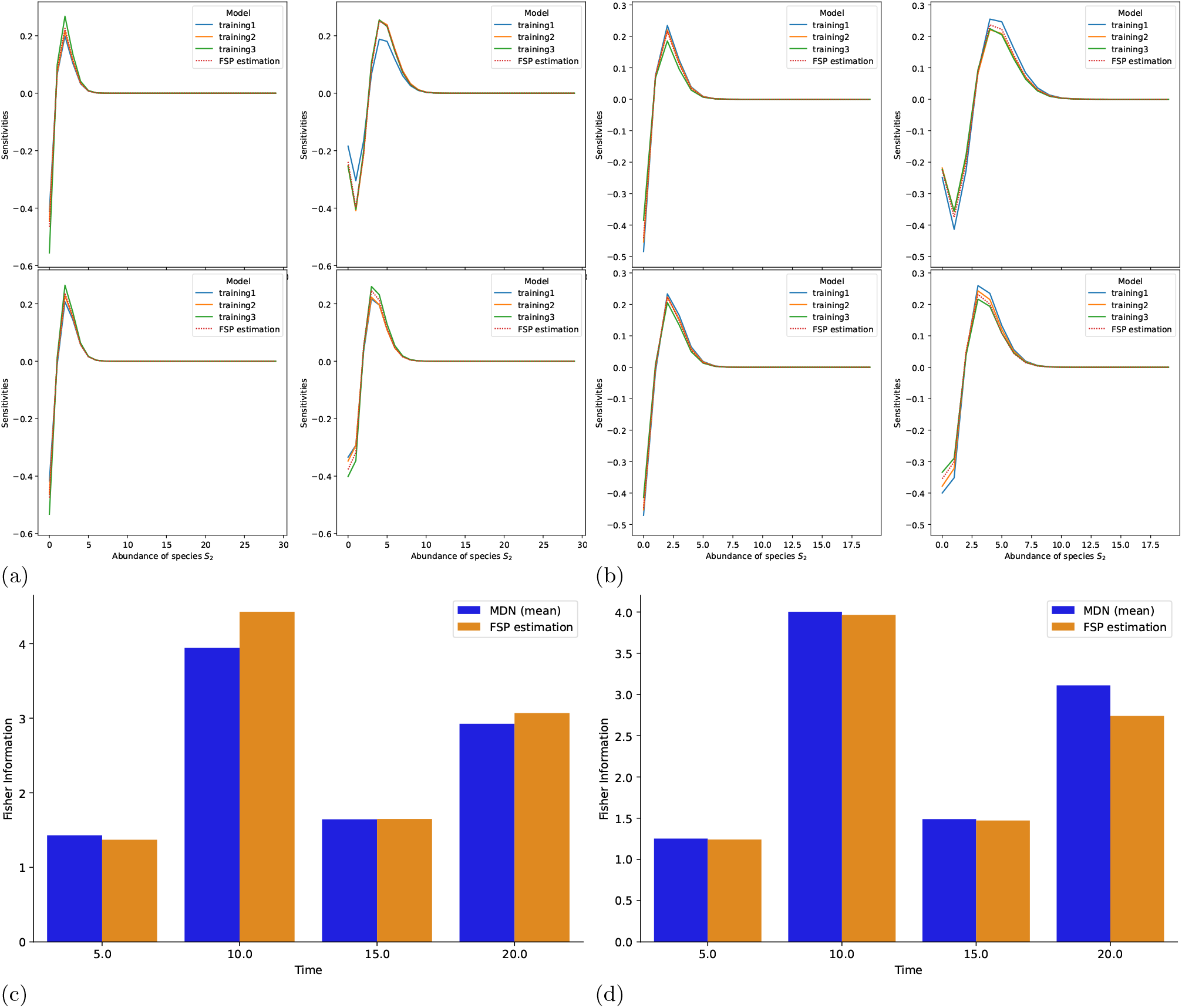
Sensitivities of the likelihood with respect to *θ*_2_ in figure 7a, and with respect to *θ*_4_ in figure 7b, as well as elements 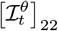 in figure 7c, and 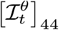 in figure 7d of the Fisher Information as a function of time for the Toggle Switch Chemical Reaction Network. Parameters: *θ*_1_ = 0.7455, *θ*_2_ = 0.3351, *θ*_3_ = 0.0078, *θ*_4_ = 0.4656, *θ*_5_ = 0.0193, *θ*_6_ = 0.2696, *θ*_7_ = 2.5266, *θ*_8_ = 0.4108, *θ*_9_ = 0.6880, *ξ*_1_ = 0.9276, *ξ*_2_ = 0.2132, *ξ*_3_ = 0.8062, *ξ*_4_ = 0.3897. For figures 7a and 7b, from left to right, top to bottom, plots correspond to times 5, 10, 15, 20.

### 3.2 Stochastic control of Chemical Reaction Networks

We now tackle the aim of controlling Chemical Reaction Networks at discrete time points, and more specifically to optimise the performance index 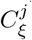 introduced in equation (3). To illustrate our approach, we deal in turn with three Chemical Reaction Networks. We introduce an alternative policy search strategy to extend the comparison with Finite State Projection-based methods.

First, we consider the controlled Production and Degradation Chemical Reaction Network specified by:

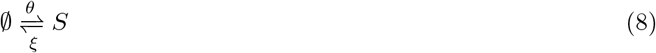

Next, we focus on the controlled Bursting Gene Chemical Reaction Network specified by:

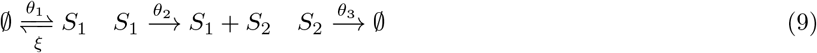

Finally, we study the Toggle Switch Chemical Reaction Network already specified by the reaction graph (7).

Given that 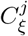 depends on the expectation of the observed species, we first evaluate the ability of our method to predict this quantity at a sequence of time points for the three controlled Chemical Reaction Networks. figure 8 illustrates the excellent accuracy of the Deep Learning-based estimation approach. In all cases, its predictions closely match the results obtained by the Finite State Projection, with a relative error below 2% throughout. Approximation of the gradient of these expectations attains comparable performance (see figures S11, S13 and S15).

**Figure 8.**
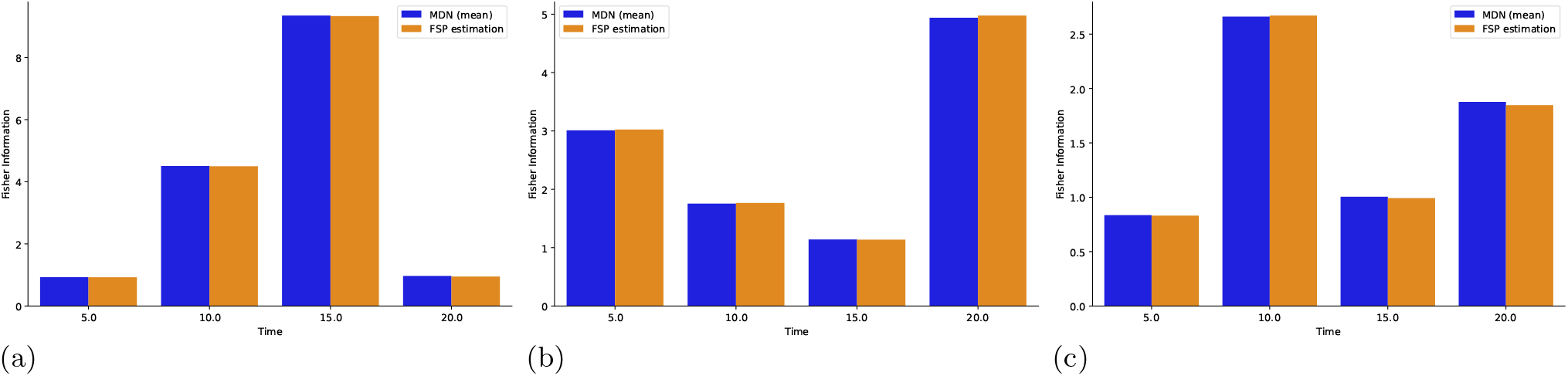
Expectation as a function of time for figure 8a the controlled Production and Degradation Chemical Reaction Network, figure 8b the controlled Bursting Gene Chemical Reaction Network and figure 8c the Toggle Switch Chemical Reaction Network. figure 8a, parameters: *θ* = 1.6869, *ξ*_1_ = 1.8234, *ξ*_2_ = 0.3082, *ξ*_3_ = 0.1011, *ξ*_4_ = 1.7771. figure 8b, parameters: *θ*_1_ = 0.4622, *θ*_2_ = 4.9699, *θ*_3_ = 0.7501, *ξ*_1_ = 0.4677, *ξ*_2_ = 1.3301, *ξ*_3_ = 2.2814, *ξ*_4_ = 0.0594. figure 8c, parameters: *θ*_1_ = 0.7455, *θ*_2_ = 0.3351, *θ*_3_ = 0.0078, *θ*_4_ = 0.4656, *θ*_5_ = 0.0193, *θ*_6_ = 0.2696, *θ*_7_ = 2.5266, *θ*_8_ = 0.4108, *θ*_9_ = 0.6880, *ξ*_1_ = 0.9276, *ξ*_2_ = 0.2132, *ξ*_3_ = 0.8062, *ξ*_4_ = 0.3897, respectively.

Having accurate estimates of the gradient of the performance index, we perform policy search to optimize 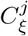 (see section S9). We evaluate the performance of the corresponding control policy by using the estimated optimal control parameter values in 10^4^ Stochastic Simulation Algorithm simulations. As shown in figures 9, 10 and 11, the results of the optimisation process are strikingly precise, even for time-varying targets.

**Figure 9.**
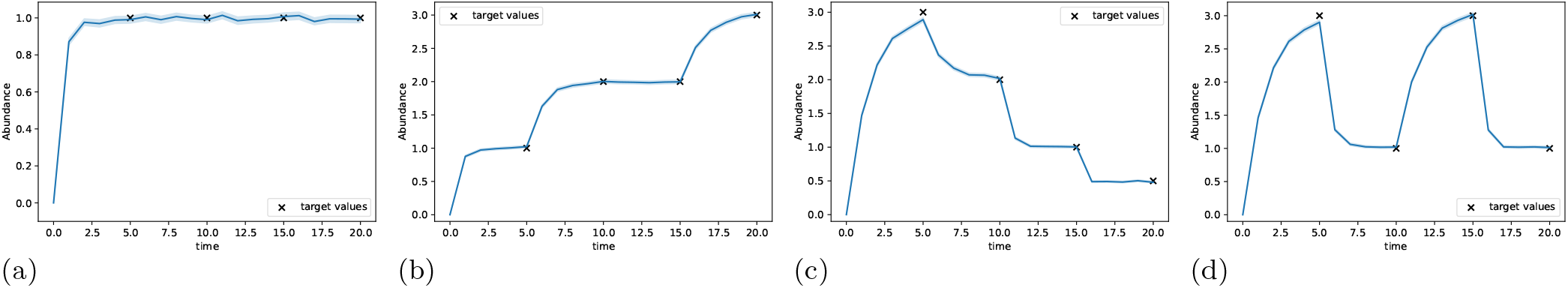
Evolution of the expected abundance of *S* for the Production and Degradation Chemical Reaction Network using the parameters *ξ*^*^ obtained by policy search. Results, parameters, tasks and hyperparameters are detailed in tables S9, S13, S14 and S15.

**Figure 10.**
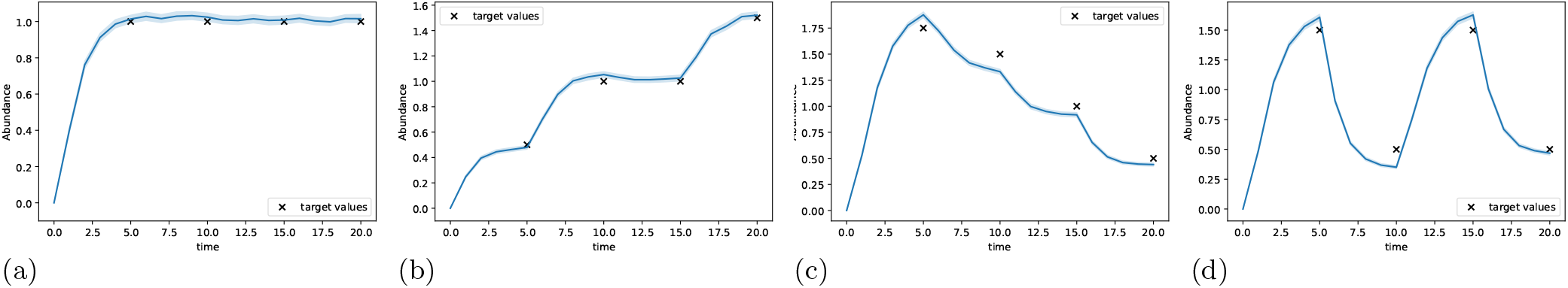
Evolution of the expected abundance of *S*_2_ for the Bursting Gene Chemical Reaction Network using the parameters *ξ*^*^ obtained by policy search. Results, parameters, tasks and hyperparameters are detailed in tables S10, S13, S16 and S17.

**Figure 11.**
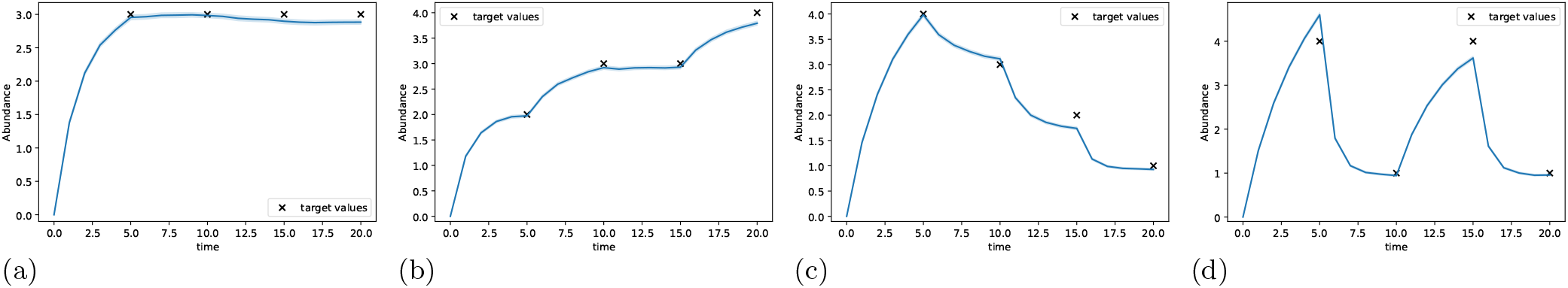
Evolution of the expected abundance of *S*_2_ for the Toggle Switch Chemical Reaction Network using the parameters *ξ*^*^ obtained by policy search. Results, parameters, tasks and hyperparameters are detailed in tables S11, S13, S18 and S19.

A crucial strength of the Deep Learning-based method is the speed at which the gradient of the likelihood can be evaluated. This results in the policy search taking no more than 7 minutes to complete for all control objectives and Chemical Reaction Networks (see subsection S13.1). To make a fair comparison with the Deep Learning method, the Finite State Projection approach was carried out in the exact same setting except for the threshold level *ε*, which was set to be the loss attained by the policy optimised using the Mixture Density Network (see table S12). The computational time for the Finite State Projection method is significantly higher than that for the Deep Learning approach, taking 31 to 824 times longer (see table S12).

Interestingly, the policy search sometimes outputs optimal parameter values outside of the training range, allowing good control performance (see tables S9, S10 and S11). For instance, in experiment 9c of the Production and Degradation Chemical Reaction Network, the value of 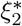 is twice the upper bound used for training. Similar observations can be made for the value of 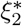 in experiment 10d of the Bursting Gene Reaction Network, as well as for the values of 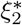 and 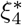 in experiment 11d of the Toggle Switch Reaction Network, which are all 1.5 times higher than the training upper bounds. This observation illustrates the practical usefulness of the generalisation ability of the Mixture Density Network.

## 4 Discussion

In this study, we used a collection of examples to demonstrate the effectiveness of our Deep Learning method in computing the Fisher Information of partially observed, sampled Stochastic Chemical Reaction Networks by comparing its results with both known ground truths and the current state-of-the-art. Results for two additional examples are included in section S11 and further support these conclusions.

The three first Chemical Reaction Networks selected here were chosen so as to parallelise the original study which introduced the Finite State Projection method for the computation of the Fisher Information [3]. For these examples, predictions by the Deep Learning method appear reliable, with a limited effect of truncation and approximation error compounding. This means that the approach could be used for Optimal Experimental Design applications similar to those showcased in the report mentioned, like the selection of optimal sampling times. The selection of species to monitor is another crucial design choice in contemporary Molecular Biology as, typically, only a limited number of species can be concurrently studied in experiments [13]. As one-dimensional marginals are directly obtained from the Deep Learning approach, the information content associated to the observation of each species can be compared and used as a principled selection criterion.

When it comes to accuracy, the Deep Learning methodology offers several benefits over the current state-of-the-art. First, no premature state-space truncation degrades the accuracy of the estimated mass function [4]. Second, the results presented here indicate that the Mixture Density Network is able to correctly predict the Fisher Information even at times further in the future than the training range. In contrast, the actual application of the Finite State Projection relies on numerical methods that have to proceed sequentially, following a time grid: getting a value at a future time requires getting them at all intermediate times as well. Still, the Deep Learning method comes with no more error guarantee than the Finite State Projection approach, and it could be less accurate in predicting the mass function and sensitivity of rarely visited states. Also, note that the Finite State Projection itself could be used to train a Neural Network outputting mixture parameters as in [6], or even directly the probability of the states in the truncated state-space. With such an additional Deep Learning layer, the Finite State Projection is itself expected to be able to generalise its predictions to future times as well.

The Fisher Information can be computed using a family of equivalent representations, and a route for further improving the Deep Learning method could thus be to judiciously select the expression for which it is most accurate (see subsection S4.2). So far, we investigated the use of two alternate formulas: one involving square roots, known as the Schrödinger kinetic energy, and a second involving second order derivatives [14, 15]. In our experience, both lead to gradient explosion.

The emergence of an expanding number of Neural Network models designed to approximate the mass function of Reaction Networks holds the possibility of enhancing the accuracy of the methodology presented here in the future. Most notably, the Variational Autoregressive Network architecture has been demonstrated to successfully estimate joint distributions, even in high-dimensional settings [8]. Still, at present, the parameters of the Reaction Network are not explicit inputs to the surrogate model, which means sensitivities cannot be obtained in the same way as in our approach. In addition, the autoregressive structure defines a family of Neural Networks trained on a time grid, which does not endow it with the ability to generalise at future times, as the Mixture Density Network can.

Standard optimisation techniques in Optimal Experimental Design commonly utilise exhaustive search, which requires to calculate quantities such as Fisher Information an extremely high number of times. The Deep Learning method seeks to strike a balance in the tradeoff between speed and accuracy. For the Chemical Reaction Networks presented above, the relative time performance of the approach improves dramatically even for limited increases in reaction graph complexity. This is unsurprising given that the Finite State Projection is well-known to suffer from the curse of dimensionality when the number of species, and hence the cardinality of the truncated state-space considered, increases. On the other hand, the time performance of the Deep Learning method will deteriorate with an increase in the initial simulation time, in particular when reactions become more frequent. At the same time, observe that simulations and training correspond to fixed costs, which means that any subsequent computation of the Fisher Information only comes at the marginal cost of the computation of the derivative by automatic differentiation, making the approach well suited for repeated evaluation of the Fisher Information.

Leveraging the estimated sensitivities a second time, we have shown how the Mixture Density Network can be used to drastically speed up policy search when compared to an approach relying exclusively on the Finite State Projection. For this second aim, it is the gradient of a performance index which needs to be repeatedly computed. The practical usefulness of the generalisation ability of the Mixture Density Network is demonstrated by the possibility given to the Projected Gradient Descent to output optimal parameters outside of the training range.

Finally, note that the time requirement for the training of the Neural Network could be further reduced by using a low-level programming language for the Stochastic Simulation Algorithm simulations, or by using approximation schemes such as the *τ* -leap algorithm [16]. A strategy to by-pass simulations altogether would be to use the same loss function as that of Physics-Informed Neural Networks, which would rely here on the discrepancy between the time derivative of the mass function as known from the Chemical Master Equation and its estimation using the Mixture Density Network [17]. This approach could only be used for single-species Chemical Reactions Networks as the Master Equation involves joint distributions, whereas the current Neural Network only outputs one-dimensional marginals. On top of this, in contrast to the approach adopted so far, this other loss function neglects the fact that the Master Equation actually deals with mass functions, likely hampering the performance of this alternative strategy.

## 5 Conclusion

The initial insight of this investigation has been to identify the potential of Mixture Density Networks in jointly estimating the mass function, sensitivities, and ultimately quantities involving both quantities. This study showed that the associated Deep Learning method can be used to compute the Fisher Information and perform policy search in an accurate and fast manner, opening an additional route for Optimal Experimental Design and Control in Molecular Biology. Future research efforts will focus on enhancing the performance of the Neural Network and extending this approach to encompass joint probabilities.

