## Supplementary Information for "Efficient Fisher Information Computation and Policy Search in Sampled Stochastic Chemical Reaction Networks through Deep Learning"

CTSB

April 2023

#### S1 Implementation of the two methods

Most numerical experiments for this paper were executed on a Mac-Book Air equipped with a 3.2 GHz Apple M1 processor and 8 Go of RAM. When a high-performance computing cluster was used, the corresponding time the script would have taken to run on 4 CPU cores of a standard private computer is still manageable.

Particular care has been devoted to take advantage of the latest advances in Deep Learning, and whenever possible, to make the Python code both time and memory efficient. For the Deep Learning-based approach:

- PyTorch has been selected as the framework to specify and train the Neural Networks [1].
- Inputs for simulations have been generated using a Low Discrepancy Sequence, the Sobol sequence [2].
- Stochastic Simulations have been parallelized across CPUs.
- Neural Network weights have been initialised using the Glorot Uniform method [3], and the learning rate has been decayed using the Cosine Annealing scheduler [4].
- Training of Neural Networks has been parallelized for hyperparameter tuning.

For the Finite State Projection-based method:

- Finite subsets of $\mathbb{N}^N$ have been enumerated in a systematic way using Cantor pairing functions [5].
- High-dimensional matrices have been stored in a sparse format.

### S2 Main notations and conventions

Given a finite set $E$, the number of elements in $E$ is denoted by $\text{Card}(E)$.

The set of integers is denoted by $\mathbb{Z}$, the set of non-negative integers $\{0, 1, \dots\}$ by $\mathbb{N}$, and the set of positive integers $\{1, 2, \dots\}$ by $\mathbb{N}^*$. Given two integers $a$ and $b$, the set of integers between $a$ and $b$ inclusive is denoted by $[\![a, b]\!]$. Given a vector $x_t \in \mathbb{N}^n$, its $i$-th element is denoted $x_t^i$.

The set of real numbers is denoted by $\mathbb{R}$, the sets of non-negative real numbers by $\mathbb{R}_+$, and the set of positive real numbers by $\mathbb{R}_+^*$.

Given a set $E$, the set of matrices of size $(N, M)$ with coefficients in $E$ is denoted by $\mathcal{M}_{N,M}(E)$. Given a vector $x$, the square matrix whose diagonal elements are the elements of $x$ is denoted by $\text{Diag}(x)$.

Given a set $E$, the space of measurable, square integrable functions defined on $E$ is denoted by $L^2(E)$.

Given a variable $\theta$ and a function $f_\theta$ of, among others, $\theta$, the gradient of $f$ with respect to $\theta$ is denoted by $\nabla_\theta f$. We choose the convention of writing gradients as row vectors. The partial derivative of $f$ with respect to the $i$-th element of $\theta$ is denoted by $\frac{\partial f}{\partial \theta_i}$.

Given a parameter $\theta$, the expectation of a random variable $X_t$ whose likelihood depends on $\theta$ is denoted by $E_\theta[X_t]$. Likewise, its variance is denoted by $V_\theta[X_t]$.

Given a set $E \subset \mathbb{R}^n$, the uniform distribution is denoted by $\mathcal{U}(E)$. The exponential distribution is denoted by $\mathcal{E}$.

### S3 Main definitions and results on Chemical Reaction Networks

Recall that we write $N \in \mathbb{N}^*$ the number of molecular species, labelled $S_i$ for $i \in [\![1, N]\!]$, and $M \in \mathbb{N}^*$ the number of chemical reactions.

Let us introduce an initial value $x_0 \in \mathbb{N}^N$, together with a stoichiometry matrix $s \in \mathcal{M}_{N,M}(\mathbb{Z})$, and a parametric propensity function $\lambda$ which, without loss of generality for both methods and except stated otherwise, we choose to be a mass-action propensity function $\lambda : \mathbb{N}^N \times \mathbb{R}_+^M \to \mathbb{R}_+^M$ defined for $j \in [\![1, M]\!]$ by:

$$\lambda_j(x_t; \theta) = \theta_j \prod_{i=1}^{N} \frac{x_t^i!}{(x_t^i - \nu_{ij})!}$$

where $\theta_j \in \mathbb{R}_+$ is a system parameter, and $\nu \in \mathcal{M}_{N,M}(\mathbb{N})$ is the reactants matrix.

The stoichiometry matrix and propensity function are jointly, implicitly defined when each reaction $j$ is given as:

$$\sum_{i=1}^{N} \nu_{ij} S_i \xrightarrow{\theta_j} \sum_{i=1}^{N} \nu'_{ij} S_i$$

where $\theta_j \in \mathbb{R}_+$ and $\nu \in \mathcal{M}_{N,M}(\mathbb{N})$ as above, and $\nu' \in \mathcal{M}_{N,M}(\mathbb{N})$ is the products matrix, such that: $s = \nu' - \nu$. Two such examples are given by reaction graphs (5) and (6).

#### S3.1 Probability mass function of Chemical Reaction Networks

The probability mass function of a Chemical Reaction Network is known to satisfy the following Partial Differential Equation, known as the Chemical Master Equation:

$$\boxed{\frac{\partial p^\theta}{\partial t} = A^\theta p^\theta} \tag{S1}$$

where $A^\theta$ is the adjoint generator of the stochastic process $(X_t)_{t\in\mathbb{R}_+}$. Together with the initial value, $A^\theta$ determines the probability mass function $p_t^\theta$ of $X_t$, and thereby fully specifies the model.

##### S3.2 Sensitivity of the likelihood of Chemical Reaction Networks

Let us define the sensitivity of the likelihood of $X_t$ in $x \in \mathbb{N}$ as:

$$\boxed{\nabla_\theta p(x; t, \theta)}$$

We set $p(-1; t, \theta) = 0$ by convention.

#### S4 General results on the Fisher Information

##### S4.1 Interpretation of the Fisher Information

The Fisher Information is rooted deep in classical Statistics, Information Geometry and Information Theory [6]. For the Production and Degradation Chemical Reaction Network, its closed-form expression bears a self-standing interpretation. First notice that, in the initial parameterization, $\mathcal{I}_t^\theta \in \mathcal{M}_{2,2}(\mathbb{R})$. The Reaction Network can be reparameterized using a lump parameter $\lambda_t^\theta$, thereby reducing its Fisher Information to the following scalar:

$$\mathcal{I}_t^{\lambda_t^\theta} = 1/\lambda_t^\theta$$

with: $\lambda_t^\theta = \theta_1/\theta_2(1-e^{-\theta_2 t})$. This allows us to visualise the Fisher Information of the Reaction Network in the plane.

Its value as a function of $\theta = (\theta_1, \theta_2)$ is represented in figure S1. Setting the production rate $\theta_1$, the Fisher Information increases with the degradation rate $\theta_2$; setting the degradation rate $\theta_2$, it decreases as the production rate $\theta_1$ increases. A Production and Degradation Reaction Network is then most informative in the relatively low production regime. A similar comment could likewise be made in the case of Pure Production (see subsection S10.3.2). By the Poisson Law of Large Numbers, we know that for large production rates, Pure Production processes essentially behave deterministically, and that their abundances grow linearly. Informally, the distinctive, stochastic behaviour of the low production regime then carries more information than the almost deterministic, linear one.

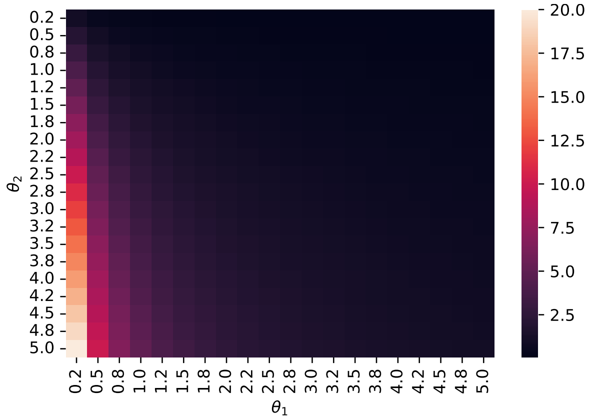

Figure S1: Dependence of the Fisher Information on the parameters of the Production and Degradation Reaction Network. Time: $t = 100$.

Also observe that as time increases, the Fisher Information does as well, and converges to $\theta_2/\theta_1$: late samples are more informative than early ones, while the maximal information content is upper bounded.

#### S4.2 Alternate definitions of the Fisher Information

Analytical calculations leading to closed-form expressions for the Fisher Information of $X_t$ leverage an explicit knowledge of $p$ and $\nabla_\theta p$. They are often made easier when starting from an alternate definition for $\mathcal{I}_t^\theta$, which is given by:

$$\boxed{\mathcal{I}_t^\theta = V_\theta\left[\nabla_\theta \log p(X_t;t,\theta)\right]}$$

*Proof.* The König-Huygens formula implies that:

$$V_\theta\left[\nabla_\theta \log p(X_t;t,\theta)\right] = E_\theta\left[\left(\nabla_\theta \log p(X_t;t,\theta)\right)^\top\left(\nabla_\theta \log p(X_t;t,\theta)\right)\right] - E_\theta\left[\nabla_\theta \log p(X_t;t,\theta)\right]^\top E_\theta\left[\nabla_\theta \log p(X_t;t,\theta)\right]$$

where:

$$\begin{aligned}
E_\theta\left[\nabla_\theta \log p(X_t;t,\theta)\right] &= \sum_{x\in\mathbb{N}^N} \nabla_\theta \log p(x;t,\theta) \times p(x;t,\theta), \text{ by the chain rule}\\
&= \sum_{x\in\mathbb{N}^N} \frac{1}{p(x;t,\theta)}\nabla_\theta p(x;t,\theta) \times p(x;t,\theta)\\
&= \sum_{x\in\mathbb{N}^N} \nabla_\theta p(x;t,\theta)\\
&= \nabla_\theta \sum_{x\in\mathbb{N}^N} p(x;t,\theta), \text{ assuming that the gradient and the sum can be permuted}\\
&= \nabla_\theta\ 1\\
&= 0
\end{aligned}$$

□

To assess whether the accuracy of the Deep Learning method could be further improved by starting from other, equivalent expressions, we leveraged two of these. The first one is the Schrödinger kinetic energy [7]:

$$\boxed{\mathcal{I}_t^\theta = 4\sum_{x\in\mathbb{N}^N}\left(\nabla_\theta\sqrt{p(x;t,\theta)}\right)^\top\left(\nabla_\theta\sqrt{p(x;t,\theta)}\right)}$$

*Proof.* In this proof, let us consider the more general case where $\alpha \in \mathbb{R} - \{-1, 1\}$ [8]. We have:

$$\begin{aligned}
\frac{4}{1-\alpha^2}\sum_{x\in\mathbb{N}^N}\frac{\partial p^{\frac{1-\alpha}{2}}}{\partial\theta_i}(x;t,\theta)\frac{\partial p^{\frac{1+\alpha}{2}}}{\partial\theta_j}(x;t,\theta) &= \frac{4}{1-\alpha^2}\sum_{x\in\mathbb{N}^N}\frac{1-\alpha}{2}\frac{\partial p}{\partial\theta_i}(x;t,\theta)p^{\frac{-1-\alpha}{2}}(x;t,\theta)\frac{1+\alpha}{2}\frac{\partial p}{\partial\theta_j}(x;t,\theta)p^{\frac{-1+\alpha}{2}}(x;t,\theta)\\
&= \sum_{x\in\mathbb{N}^N}\frac{1}{p(x;t,\theta)}\frac{\partial p}{\partial\theta_i}(x;t,\theta)\frac{\partial p}{\partial\theta_j}(x;t,\theta)\\
&= \sum_{x\in\mathbb{N}^N}\frac{1}{p(x;t,\theta)}\frac{\partial p}{\partial\theta_i}(x;t,\theta)\frac{1}{p(x;t,\theta)}\frac{\partial p}{\partial\theta_j}(x;t,\theta)p(x;t,\theta)\\
&= \left[\mathcal{I}_t^\theta\right]_{ij}
\end{aligned}$$

Taking $\alpha = 0$ in the first expression, we get a representation involving square roots, the Schrödinger kinetic energy:

$$\mathcal{I}_t^\theta = 4\sum_{x\in\mathbb{N}^N}\left(\nabla_\theta\sqrt{p(x;t,\theta)}\right)^\top\left(\nabla_\theta\sqrt{p(x;t,\theta)}\right)$$

□

The second expression investigated is [8]:

$$\boxed{\mathcal{I}_t^\theta = -E_\theta\left[\nabla_\theta^2 \log p(X_t;t,\theta)\right]}$$

*Proof.*

$$
\begin{aligned}
-\sum_{x\in\mathbb{N}^N} \frac{\partial^2 \log p}{\partial\theta_i \partial\theta_j}(x;t,\theta)p(x;t,\theta) &= -\sum_{x\in\mathbb{N}^N} \frac{\partial}{\partial\theta_i}\left(\frac{\partial \log p}{\partial\theta_j}\right)(x;t,\theta)p(x;t,\theta) \\
&= -\sum_{x\in\mathbb{N}^N} \frac{\partial}{\partial\theta_i}\left(\frac{1}{p}\frac{\partial p}{\partial\theta_j}\right)(x;t,\theta)p(x;t,\theta) \\
&= -\sum_{x\in\mathbb{N}^N} \left(\frac{-1}{\big(p(x;t,\theta)\big)^2}\frac{\partial p}{\partial\theta_i}(x;t,\theta)\frac{\partial p}{\partial\theta_j}(x;t,\theta) + \frac{1}{p(x;t,\theta)}\frac{\partial^2 p}{\partial\theta_i\partial\theta_j}(x;t,\theta)\right)p(x;t,\theta) \\
&= \sum_{x\in\mathbb{N}^N} \frac{1}{p(x;t,\theta)}\frac{\partial p}{\partial\theta_i}(x;t,\theta)\frac{\partial p}{\partial\theta_j}(x;t,\theta) + \sum_{x\in\mathbb{N}^N}\frac{\partial^2 p}{\partial\theta_i\partial\theta_j}(x;t,\theta) \\
&= \left[\mathcal{I}_t^\theta\right]_{ij} + \frac{\partial^2}{\partial\theta_i\partial\theta_j}\sum_{x\in\mathbb{N}^N} p(x;t,\theta) \\
&= \left[\mathcal{I}_t^\theta\right]_{ij}
\end{aligned}
$$

□

### S5 Deep Learning approximation of the likelihood and its sensitivities

#### S5.1 Estimated likelihood

##### S5.1.1 Activation Functions and element-wise Reference Functions

Before introducing the architecture of the Neural Network, let us first define the following Activation Functions:

- Softmax Activation Function

$$
\begin{aligned}
\text{Softmax}: \quad & \mathbb{R}^n \longrightarrow [0,1]^n \\
& x = (x_1, ..., x_n) \mapsto \left(\frac{\exp(x_i)}{\sum_{j=1}^{n}\exp(x_j)}\right)_{1\le i\le n}
\end{aligned}
$$

- ReLU Activation Function

$$
\begin{aligned}
\text{ReLU}: \quad & \mathbb{R}^n \longrightarrow \mathbb{R}_+^n \\
& x = (x_1, ..., x_n) \mapsto \big(\max(x_i, 0)\big)_{1\le i\le n}
\end{aligned}
$$

- Sigmoid Activation Function

$$
\begin{aligned}
\sigma: \quad & \mathbb{R}^n \longrightarrow [0,1]^n \\
& x = (x_1, ..., x_n) \mapsto \left(\frac{1}{1+\exp(-x_i)}\right)_{1\le i\le n}
\end{aligned}
$$

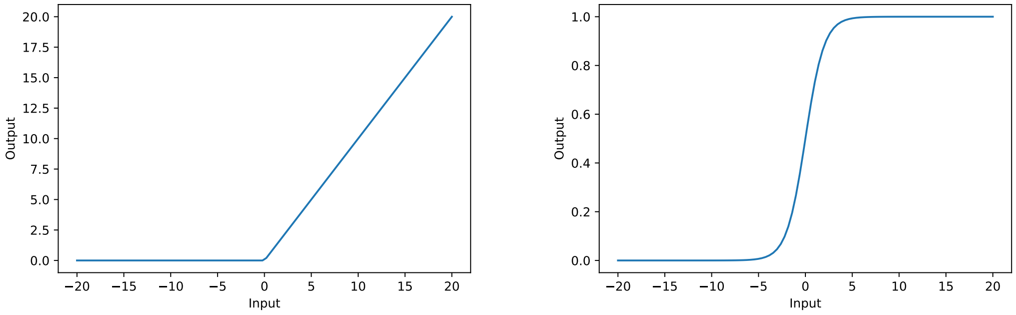

(a) Graph of the ReLU function

(b) Graph of the Sigmoid function

Figure S2: Graphs of Activation Functions when the input $x$ is in $\mathbb{R}$

Let us also define the following element-wise Reference Functions:

- Element-wise logarithm

$$\begin{array}{rccc} \log_{10}: & (\mathbb{R}_+^*)^n & \longrightarrow & \mathbb{R}^n \\ & x = (x_1, ..., x_n) & \mapsto & \Big(\log_{10}(x_i)\Big)_{1 \le i \le n} \end{array}$$

- For $A \subset \mathbb{R}^n$, element-wise indicator

$$\begin{array}{rccc} \mathbb{1}_A: & \mathbb{R}^n & \longrightarrow & \mathbb{R}^n \\ & x = (x_1, ..., x_n) & \mapsto & \big(\mathbb{1}_A(x_i)\big)_{1 \le i \le n} \end{array}$$

##### S5.1.2 Architecture of the Mixture Density Network

Let us introduce $K \in \mathbb{N}^*$. The Mixture Density Network is a function NN of the form:

$$\begin{array}{rccc} \mathrm{NN} = \big(\mathrm{NN}_1, \mathrm{NN}_2, \mathrm{NN}_3\big): & \mathbb{R}_+^* \times (\mathbb{R}_+^*)^M & \longrightarrow & [0,1]^K \times \mathbb{R}_+^K \times [0,1]^K \\ & (t, \theta) & \mapsto & (w, r, q) \end{array}$$

It takes as inputs:

- $t \in \mathbb{R}_+^*$, the time.
- $\theta \in (\mathbb{R}_+^*)^M$, the vector of system parameters of the chosen Chemical Reaction Network.

Following [9], we start by taking the $\log_{10}$ of $(t, \theta)$, which we use to define the input layer. In our experience, this initial transformation speeds up training.

To specify the outputs, let us first define:

- $A_h \in \mathcal{M}_{N_{\text{hidden}}, 1+M}(\mathbb{R})$ the matrix of the weights of the hidden layer.
- $B_h \in \mathbb{R}^{1+M}$ the vector of its biases.
- $A_{o1}, A_{o2}, A_{o3} \in \mathcal{M}_{K, N_{\text{hidden}}}(\mathbb{R})$ the matrix of the weights of the $1^{st}$, $2^{nd}$ and $3^{rd}$ output layers.
- $B_{o1}, B_{o2}, B_{o3} \in \mathbb{R}^K$ the vectors of their biases.

The Mixture Density Network returns as outputs:

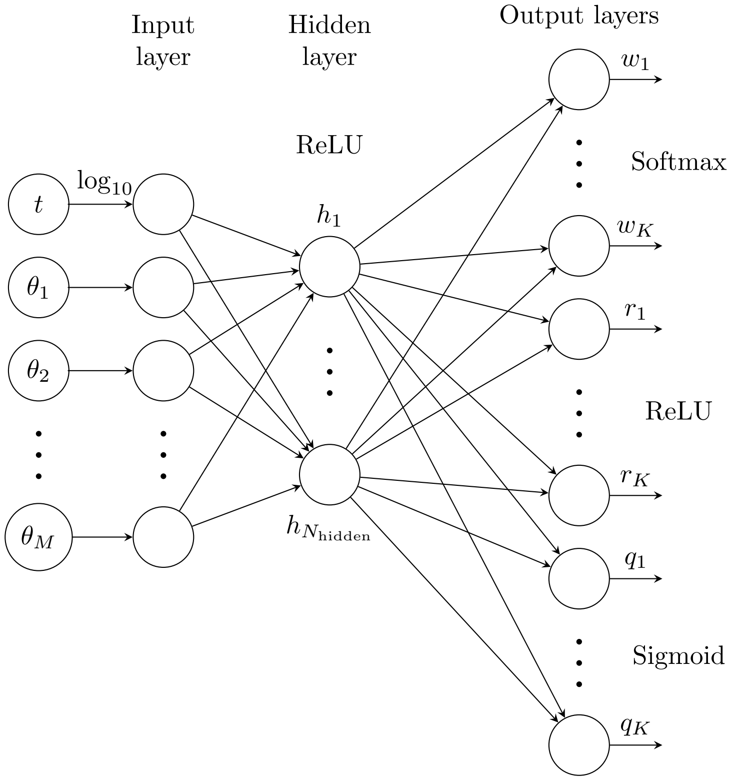

Figure S3: Architecture of the Mixture Density Network

- $w \in [0,1]^K$ such that:

$$\forall k \in [\![1, K]\!], \quad \boxed{w_k(h) = \Big[\mathrm{Softmax}(A_{o1}h + B_{o1})\Big]_k}$$

- $r \in \mathbb{R}_+^K$ such that:

$$\forall k \in [\![1, K]\!], \quad r_k(h) = \mathrm{ReLU}\Big([A_{o2}h + B_{o2}]_k\Big)$$

$$\boxed{r_k(h) = \mathrm{ReLU}\Big(\sum_{j=1}^{N_{\text{hidden}}} [A_{o2}]_{kj} h_j + [B_{o2}]_k\Big)}$$

- $q \in [0,1]^K$ such that:

$$\forall k \in [\![1, K]\!], \quad q_k(h) = \sigma\Big([A_{o3}h + B_{o3}]_k\Big)$$

$$\boxed{q_k(h) = \sigma\Big(\sum_{j=1}^{N_{\text{hidden}}} [A_{o3}]_{kj} h_j + [B_{o3}]_k\Big)}$$

where $h \in \mathbb{R}_+^{N_{\text{hidden}}}$ is the vector of hidden layer outputs, defined as:

$$\forall j \in [\![1, N_{\text{hidden}}]\!], \quad h_j(t, \theta) = \mathrm{ReLU}\Big([A_h]_{j1} \log_{10}(t) + \sum_{i=1}^{M} [A_h]_{j,i+1} \log_{10}(\theta_i) + [B_h]_j\Big)$$

The outputs of the Neural Network are used as $K$ sets of mixture parameters to define a Negative Binomial Mixture Model $\hat{p} : \mathbb{N}^N \times \mathbb{R}_+ \times \mathbb{R}_+^M \to \mathbb{R}_+$ as:

$$\boxed{\hat{p}(x; t, \theta) = \sum_{k=1}^{K} \mathrm{NN}_1^k(t, \theta) p_{\mathrm{NB}}\big(x; \mathrm{NN}_2^k(t, \theta), \mathrm{NN}_3^k(t, \theta)\big)}$$

Note that with only slight modifications to its architecture, the Neural Network can output mixture parameters of other distributions, like the Poisson distribution.

In the expression above, also observe that $\mathrm{NN}_2^k$ takes values in $\mathbb{R}_+$. In section S10.1.3, the distribution $p_{\mathrm{NB}}$ of a Negative Binomial random variable is specified by two parameters, $r \in \mathbb{N}^*$, and $q \in (0, 1]$. The definition of Negative Binomials can be extended by first generalising factorials using the Gamma function $\Gamma : x \in \mathbb{R}_+^* \mapsto \int_0^\infty t^{x-1} e^{-t} dt$. We next define binomial coefficients as:

$$\binom{b}{a} = \begin{cases} \frac{\Gamma(b+1)}{\Gamma(a+1)\Gamma(b-a+1)} & \text{if } (a, b) \in (-1, +\infty)^2 \\ 1 & \text{if } a = -1, b = -1 \\ 0 & \text{if } a = -1, b \in (-1, +\infty) \end{cases}$$

This generalisation allows in turn to extend the definition of the Negative Binomial distribution to $r$ in $\mathbb{R}_+$:

$$\forall q \in (0, 1], \quad p_{\mathrm{NB}}(k; r, q) = \begin{cases} \frac{\Gamma(k+r)}{\Gamma(r)\Gamma(k+1)} (1 - q)^k q^r & \text{if } (k, r) \in \mathbb{N} \times \mathbb{R}_+^* \\ 1 & \text{if } (k, r) = (0, 0) \\ 0 & \text{if } (k, r) \in \mathbb{N}^* \times \{0\} \end{cases}$$

##### S5.1.3 Datasets generation

Subsequent paragraphs will repeatedly refer to training, validation and testing datasets. Their usage is detailed below, and only their generation is covered here. Parameter configurations $\theta$ of a Chemical Reaction Network are sampled over finite intervals using a Low Discrepancy Sequence, the Sobol sequence. The corresponding datasets $\mathcal{D}_t^\theta$ are then built using the Stochastic Simulation Algorithm [10], and used to generate estimated distributions of $X_t$, which are jointly stored in a dataset $\mathcal{D}$. The training, validation and testing datasets are generated by splitting this initial dataset into smaller ones. This ensures in particular that the training and testing datasets do not overlap.

##### S5.1.4 Network fitting

In our method, the optimisation of one Neural Network requires the samples from one training dataset. We choose the Küllback-Leibler Divergence loss as the performance metric for our training procedure. For a predicted vector $\hat{y} \in [0, 1]^n$ and a target vector $y \in [0, 1]^n$, the Küllback-Leibler Divergence loss is given by:

$$\boxed{\mathrm{KL}(y, \hat{y}) = \sum_{i=1}^{n} y_i \log\left(\frac{y_i}{\hat{y}_i}\right)}$$

Note that the expression above is not defined whenever $y_i = 0$ for some $i \in [\![1, n]\!]$. If $y_i = 0$, whatever $\hat{y} \in [0, 1]^n$, we set: $y_i \log(y_i/\hat{y}_i) = 0$. The formula is not defined either whenever $\hat{y}_i = 0$ for some $i \in [\![1, n]\!]$. For this case as well, a convention could be adopted, but at least with PyTorch, in our experience, the associated gradients then explode. If $\hat{y}_i = 0$, we set: $y_i \log(y_i/\hat{y}_i) = y_i \log(10^{10} y_i)$.

Although the considerations above are enough to make the Küllback-Leibler loss fully defined, we choose to replace $y_i$, $\hat{y}_i \leq 10^{-10}$ by $10^{-10}$, as we observed this convention speeds up computation time. When $(y_i, \hat{y}_i) = (0, 0)$, and $y_i \neq 0$ with $\hat{y}_i = 0$, this computational choice does not contradict the mathematical convention from the previous paragraph. When $y_i = 0$ with $\hat{y}_i \leq 10^{-10}$, PyTorch sets: $y_i \log(y_i/\hat{y}_i) = 0$.

The output of the Deep Learning approach is a full-fledged probability distribution $\hat{p}$, which each time needs to be truncated to $\hat{y} \in [0,1]^n$ to compute the Küllback-Leibler loss. We choose $n$ to be the size of $y$.

Before the training, the weights of the Neural Network are initialised using the Glorot Uniform method [3]. This approach sets the weights of each layer to values sampled from a uniform distribution $\mathcal{U}(-a, a)$. In our cases, $a$ is given by:

$$\boxed{a = \begin{cases} \sqrt{\frac{6}{M+N_{\text{hidden}}}} & \text{for the hidden layer.} \\ \sqrt{\frac{6}{N_{\text{hidden}}+K}} & \text{for the } i\text{-th output layer}, i \in \{1,2,3\}. \end{cases}}$$

The chosen training dataset is used to perform optimisation iteratively, and each iteration $n$ of the procedure is referred to as an epoch. We use Adam, a highly efficient derivative of Mini-Batch Stochastic Gradient Descent, as the training routine [11], and set the maximal number of epochs to $n_{\text{epochs}}$ (see algorithm (S1)). Recall that in vanilla Stochastic Gradient Descent, each sample from the dataset is processed individually: the loss between a single prediction and target value is computed, and the corresponding gradient is used to update the weights of the Neural Network. In Mini-Batch Stochastic Gradient Descent, at each epoch, all samples are randomly allocated to batches of size $n_{\text{batches}}$: the loss is computed for each batch, and the corresponding gradients are used in turn to update the weights of the Neural Network.

**Algorithm S1** Mini-Batch Stochastic Gradient Descent

**Input:** $n_{\text{epochs}}$, $n_{\text{batches}}$, $l_{r,0}$, $\mathcal{D}_{\text{train}}$,

$A \leftarrow \mathcal{U}(-a,a)$, $B \leftarrow \mathcal{U}(-a,a)$ ▷ $a$ as defined by the Glorot Uniform method. $n_{\text{batches}}$.
$l_r \leftarrow l_{r,0}$
**for** $n \in [\![1, n_{epochs}]\!]$ **do**:
  Split $\mathcal{D}_{\text{train}}$ in shuffled batches of size
  **for** $b = \{b_1, \cdots, b_{n_{\text{batches}}}\} \in$ batches **do**:
    loss $\leftarrow \frac{1}{\text{Card}(b)} \sum\limits_{(t,\theta,\hat{y}) \in b} \text{KL}\Big(\big\{\hat{p}(k;t,\theta)\big\}_k, \hat{y}\Big)$
    $A \leftarrow A - l_r \nabla_A$loss
    $B \leftarrow B - l_r \nabla_B$loss
  **end for**
  $l_r \leftarrow l_{r,n}$ ▷ $l_{r,n}$ as defined by the Cosine Annealing scheduler
**end for**

The step size of each descent is parameterised by a learning rate, which we decay following the Cosine Annealing scheduler, as it performed best in our experience [4]. At epoch $n$, the learning rate $l_{r,n}$ used for all descents is:

$$\boxed{l_{r,n} = \begin{cases} \frac{l_{r,0}}{2}\Big(1 + \cos\big(\frac{n}{n_{\text{epochs}}}\pi\big)\Big) & \text{if } \frac{n}{n_{\text{epochs}}} \notin 2\mathbb{N}+1 \\ \frac{l_{r,0}}{2}\Big(1 - \cos\big(\frac{1}{n_{\text{epochs}}}\pi\big)\Big) & \text{else.} \end{cases}}$$

where $l_{r,0} \in \mathbb{R}_+^*$. Observe that, starting from $l_{r,0}$, the learning rate progressively decreases towards 0.

##### S5.1.5 Network testing

Once the Neural Network has been fitted, its performance should be assessed on a dataset which has not yet been used for training, called the testing dataset. This way, we assess the ability of the Neural Network to generalise its learning to data never encountered before, and to confirm that it did not overfit to data seen in the past.

Here, for a given Chemical Reaction Network, a Mixture Density Network was trained on three disjoint training datasets, and tested on a common testing dataset. In all figures, plots correspond to predictions made on samples arbitrarily taken from the middle of the testing dataset, either as three separate predictions or as their mean.

##### S5.1.6 Hyperparameter tuning

Hyperparameters significantly contribute to the performance of Neural Networks. Those involved in the specification and training of our Mixture Density Network are listed below:

| Step of the procedure | Denomination | Notation |
|---|---|---|
| Datasets generation | Number of samples in dataset | $n_{\text{sampled}}$ |
| | Number of simulations per parameter configuration | $n_{\text{sim}}$ |
| Architecture of the Neural Network | Number of mixture components | $K$ |
| | Number of neurons for the hidden layer | $N_{\text{hidden}}$ |
| Optimisation of the Neural Network | Batch size | $n_{\text{batches}}$ |
| | Initial learning rate | $l_{r,0}$ |
| | Maximum number of training epochs | $n_{\text{epochs}}$ |
| | Patience level for early stopping | $n_p$ |
| | Tolerance threshold for early stopping | $\delta$ |

Hyperparameter tuning refers to the process of choosing hyperparameters which improve the performance of the Neural Network. Here, whatever the number of Neural Networks trained, this process was carried using only one of the so-called validation datasets.

The most straightforward way to optimise the hyperparameters is to perform grid search. Following [9], in the specific context of hyperparameter tuning, we use the Hellinger Distance loss as the performance metric. For a predicted vector $\hat{y} \in [0,1]^n$ and a target vector $y \in [0,1]^n$, the Hellinger Distance loss is given by:

$$\boxed{H(y,\hat{y}) = \sqrt{1 - \sum_{k=1}^{n} \sqrt{y_i \hat{y}_i}}}$$

*Proof.* Starting from the usual definition of the Hellinger Distance between distributions $y$ and $\hat{y}$ with a support of size $n \in \mathbb{N}^*$:

$$\begin{aligned}
H(y,\hat{y}) &= \frac{1}{\sqrt{2}} \sqrt{\sum_{i=1}^{n} (\sqrt{y_i} - \sqrt{\hat{y}_i})^2} \\
&= \sqrt{\frac{1}{2} \sum_{i=1}^{n} (y_i - 2\sqrt{y_i \hat{y}_i} + \hat{y}_i)} \\
&= \sqrt{\frac{1}{2} \Big( \sum_{i=1}^{n} y_i + \sum_{i=1}^{n} \hat{y}_i \Big) - \sum_{i=1}^{n} \sqrt{y_i \hat{y}_i}} \\
&= \sqrt{1 - \sum_{k=1}^{n} \sqrt{y_i \hat{y}_i}}
\end{aligned}$$

□

Similarly as before, to compute the Hellinger Distance loss, we truncate $\hat{p}_t^{\theta}$ to a vector $\hat{y}$ to match to the size of $y$.

Direct interpretation of losses is often hard, and visualisation of the estimated distributions complements this metric-based approach by providing additional insights.

#### S5.2 Estimated sensitivities of likelihood

Our method crucially relies on the ease and efficiency of taking the gradient of a Neural Network with respect to its input. This favourable property of Neural Networks is underpinned by their compositional structure. Indeed, for a composite function, the gradient reduces to a product of elementary gradients.

To make this point explicit, for any $(t, \theta) \in \mathbb{R}_+^* \times (\mathbb{R}_+^*)^M$, let us explicitly write $w \in [0, 1]^K$, the first output of our Mixture Density Network, as $\text{NN}_1(t, \theta)$. Given the above definitions, we have:

$$\text{NN}_1(t, \theta) = \text{Softmax}\Big(A_{o1}\Big(\text{ReLU}\big(A_h \log_{10}(t, \theta) + B_h\big)\Big) + B_{o1}\Big)$$

The gradient of $\text{NN}_1$ can then be simply computed using the chain rule as:

$$\nabla_\theta \text{NN}_1 = \nabla_{x_5}\text{Softmax}\ \nabla_{x_4}(A_{o1} \cdot + B_{o1})\nabla_{x_3}\text{ReLU}\ \nabla_{x_2}(A_h \cdot + B_h)\nabla_\theta \log_{10}$$

This point is further illustrated in table S1 for all outputs.

| Forward propagation | | | Backward propagation | | |
|---|---|---|---|---|---|
| $x_1 = (t, \theta)$ | | | Id | | |
| *Input layer:* $x_2 = \log_{10}(x_1)$ | | | *Input layer:* $\nabla_{x_1} x_2 = \text{Diag}\big(\ln(10)x_1\big)^{-1}$ | | |
| *Hidden layer:* $x_3 = A_h x_2 + B_h$, $x_4 = \text{ReLU}(x_3)$ | | | *Hidden layer:* $\nabla_{x_2} x_3 = A_h$, $\nabla_{x_3} x_4 = \text{Diag}\big(\mathbb{1}_{\mathbb{R}_+^*}(x_3)\big)$ | | |
| *Output layer 1:* $x_5 = A_{o1}x_4 + B_{o1}$, $x_6 = S(x_5)$ | *Output layer 2:* $y_5 = A_{o2}x_4 + B_{o2}$, $y_6 = \text{ReLU}(y_5)$ | *Output layer 3:* $z_5 = A_{o3}x_4 + B_{o3}$, $z_6 = \sigma(z_5)$ | *Output layer 1:* $\nabla_{x_4} x_5 = A_{o1}$, $\nabla_{x_5} x_6 = \text{Diag}\big(\text{S}(x_5)\big) - S(x_5)S(x_5)^\top$ | *Output layer 2:* $\nabla_{x_4} y_5 = A_{o2}$, $\nabla_{y_5} y_6 = \text{Diag}\big(\mathbb{1}_{\mathbb{R}_+^*}(y_5)\big)$ | *Output layer 3:* $\nabla_{x_4} z_5 = A_{o3}$, $\nabla_{z_5} z_6 = \text{Diag}\big(e^{-z_{5i}}\sigma(z_5)_i^2\big)$ |

Table S1: Forward and backward propagations to illustrate automatic differentiation. For ease of reading, we called the Softmax function $S$ in the table, and with a slight abuse of notations, we assimilate here functions and their image.

The very specific structure of the gradient of Neural Networks has made it possible to make the whole differentiation fully automated in a process referred to as automatic differentiation. In the background, high-performance frameworks like PyTorch store the relation between gradients in what is called a computational graph, whose hierarchical structure for our Mixture Density Network parallels that of table S1.

Now to compute the Fisher Information, we need to obtain the gradient of the estimated likelihood $\hat{p}_t$ with respect to the inputs $\theta$. To fully illustrate that this problem is made easy when the approximator is a Neural Network, and that it can be efficiently addressed by automatic differentiation, let us give an analytical formula for this gradient. The detailed explicit expression is notation-heavy and hence hard to read. We have therefore chosen to highlight its component parts rather than its detailed expression as seen below:

$$\begin{aligned}\nabla_\theta \hat{p}(x; t, \theta) = \sum_{k=1}^{K} \Big[ &\nabla_\theta \text{NN}_1^k(t, \theta)\ p_{\text{NB}}\big(x; \text{NN}_2^k(t, \theta), \text{NN}_3^k(t, \theta)\big) \\ &+ \text{NN}_1^k(t, \theta)\Big(\nabla_{\text{NN}_2^k} p_{\text{NB}}\big(x; \text{NN}_2^k(t, \theta), \text{NN}_3^k(t, \theta)\big)\ \nabla_\theta \text{NN}_2^k(t, \theta) \\ &+ \nabla_{\text{NN}_3^k} p_{\text{NB}}\big(x; \text{NN}_2^k(t, \theta), \text{NN}_3^k(t, \theta)\big)\ \nabla_\theta \text{NN}_3^k(t, \theta)\Big)\Big]\end{aligned}$$

*Proof.*

$\forall (t, \theta) \in \mathbb{R}_+^* \times (\mathbb{R}_+^*)^M, \forall x \in \mathbb{N}^N,$

$$\begin{aligned}\nabla_\theta \hat{p}(x; t, \theta) &= \sum_{k=1}^{K} \Big[\nabla_\theta \text{NN}_1^k(t, \theta)\ p_{\text{NB}}\big(x; \text{NN}_2^k(t, \theta), \text{NN}_3^k(t, \theta)\big) + \text{NN}_1^k(t, \theta)\ \nabla_\theta p_{\text{NB}}\big(x; \text{NN}_2^k(t, \theta), \text{NN}_3^k(t, \theta)\big)\Big] \\ &= \sum_{k=1}^{K} \Big[\nabla_\theta \text{NN}_1^k(t, \theta)\ p_{\text{NB}}\big(x; \text{NN}_2^k(t, \theta), \text{NN}_3^k(t, \theta)\big) + \text{NN}_1^k(t, \theta)\Big(\nabla_{\text{NN}_2^k} p_{\text{NB}}\big(x; \text{NN}_2^k(t, \theta), \text{NN}_3^k(t, \theta)\big)\ \nabla_\theta \text{NN}_2^k(t, \theta) \\ &\quad + \nabla_{\text{NN}_3^k} p_{\text{NB}}\big(x; \text{NN}_2^k(t, \theta), \text{NN}_3^k(t, \theta)\big)\ \nabla_\theta \text{NN}_3^k(t, \theta)\Big)\Big], \text{ by the chain rule}\end{aligned}$$

$\square$

$\nabla_\theta \mathrm{NN}_1(t,\theta)$, $\nabla_\theta \mathrm{NN}_2(t,\theta)$ and $\nabla_\theta \mathrm{NN}_3(t,\theta)$ can themselves be computed using the chain rule, as illustrated in table S1.

### S6 Finite State Projection approximation of the mass function and the sensitivities of the likelihood

#### S6.1 Estimated probability mass function

Following [12], we truncate the state-space of the Chemical Reaction Network of interest to a finite subset of $\mathbb{N}^N$ with $N_{\max}$ elements, and augment it with an absorbing state. The mass function $\hat{p}^\theta$ of the non-absorbing states satisfies the following finite-dimensional Chemical Master Equation:

$$\boxed{\frac{\partial \hat{p}^\theta}{\partial t} = \hat{A}^\theta \hat{p}^\theta} \tag{S2}$$

where $\hat{A}^\theta$ is the adjoint generator of the Finite State Projection process, excluding the absorbing state. From now on, in this section, we only consider the newly defined process.

The generalised Cantor pairing function $\Phi_N$ offers a way to enumerate elements of $\mathbb{N}^N$ using elements of $\mathbb{N}$ [5]. It is defined as:

$$\forall n > 2, \quad \Phi_n : \begin{array}{ccc} \mathbb{N}^n & \longrightarrow & \mathbb{N} \\ (x_1, ..., x_n) & \mapsto & \Phi_2\big(\Phi_{n-1}(x_1, ..., x_{n-1}), x_n\big) \end{array}$$

$$\text{where} \quad \Phi_2 : \begin{array}{ccc} \mathbb{N}^2 & \longrightarrow & \mathbb{N} \\ (x_1, x_2) & \mapsto & \frac{1}{2}(x_1+x_2)(x_1+x_2+1)+x_2 \end{array}$$

In order to implement the method in a systematic way, we use this pairing function to enumerate elements of the finite state-space. More specifically, we choose a $C_r \in \mathbb{N}$ such that the $N_{\max}$-th element of the enumerated state-space corresponds to $(0, ..., 0, C_r) \in \mathbb{N}^N$. The enumerated state-space is then given by:

$$\Phi_N^{-1}\big(\llbracket \Phi_N(0, ..., 0), \Phi_N(0, ..., 0, C_r) \rrbracket\big)$$

It should be noted that $\Phi_N(0, ..., 0, C_r) = N_{\max} - 1$. In the case of $N = 2$, this means that:

$$N_{\max} = \frac{C_r(C_r+3)}{2} + 1$$

#### S6.2 Estimated sensitivities of the likelihood

The mass function $\hat{p}^\theta$ and sensitivity $\hat{S}^\theta$ of the Finite State Projection process solve the equation given by:

$$\boxed{\frac{\partial}{\partial t}\begin{pmatrix} \hat{p}^\theta \\ [\hat{S}^\theta]_{.i} \end{pmatrix} = \begin{pmatrix} \hat{A}^\theta & 0 \\ \frac{\partial \hat{A}^\theta}{\partial \theta_i} & \hat{A}^\theta \end{pmatrix} \begin{pmatrix} \hat{p}^\theta \\ [\hat{S}^\theta]_{.i} \end{pmatrix} \quad \forall i \in \llbracket 1, M \rrbracket} \tag{S3}$$

where $\hat{A}^\theta \in \mathcal{M}_{N_{\max}, N_{\max}}(\mathbb{R})$, and with $[\hat{S}_0^\theta]_{.i} = 0$, as the initial distribution is not parameterized by $\theta$.

*Proof.* The finite-dimensional Chemical Master Equation (S2) can be differentiated as follows:

$$\forall i \in [\![1, M]\!], \frac{\partial}{\partial \theta_i}\Big[\frac{\partial \hat{p}}{\partial t}(\cdot; t, \theta) = \hat{A}^\theta \hat{p}(\cdot; t, \theta)\Big]$$

$$\frac{\partial}{\partial \theta_i}\Big[\frac{\partial \hat{p}}{\partial t}(\cdot; t, \theta)\Big] = \frac{\partial \hat{A}^\theta}{\partial \theta_i}\hat{p}(\cdot; t, \theta) + \hat{A}^\theta \frac{\partial \hat{p}}{\partial \theta_i}(\cdot; t, \theta) \text{ by the product rule}$$

Assuming the two derivatives can be permuted, we then have:

$$\forall i \in [\![1, M]\!], \frac{\partial}{\partial t}\Big[\frac{\partial \hat{p}}{\partial \theta_i}(\cdot; t, \theta)\Big] = \frac{\partial \hat{A}^\theta}{\partial \theta_i}\hat{p}(\cdot; t, \theta) + \hat{A}^\theta \frac{\partial \hat{p}}{\partial \theta_i}(\cdot; t, \theta)$$

$$\frac{\partial [\hat{S}]_{.i}}{\partial t}(\cdot; t, \theta) = \frac{\partial \hat{A}^\theta}{\partial \theta_i}\hat{p}(\cdot; t, \theta) + \hat{A}^\theta \big[\hat{S}\big]_{.i}(\cdot; t, \theta) \tag{S4}$$

Equations (S2) and (S4) jointly give the desired the result. □

Introduce $\alpha \in \mathbb{N}^*$, and let $I = \{i_1, i_2, ..., i_\alpha\} \subset [\![1, M]\!]$ be a set of parameter indices. The sensitivity of the likelihood with respect to multiple parameters can be obtained at once by solving:

$$\frac{\partial}{\partial t}\begin{pmatrix} \hat{p}^\theta \\ [\hat{S}^\theta]_{.i_1} \\ \vdots \\ [\hat{S}^\theta]_{.i_\alpha} \end{pmatrix} = \begin{pmatrix} \hat{A}^\theta & 0 & 0 & \dots & 0 \\ \frac{\partial \hat{A}^\theta}{\partial \theta_{i_1}} & A & 0 & \dots & 0 \\ \vdots & \vdots & \vdots & \ddots & \vdots \\ \frac{\partial \hat{A}^\theta}{\partial \theta_{i_\alpha}} & 0 & 0 & \dots & A \end{pmatrix}\begin{pmatrix} \hat{p}^\theta \\ [\hat{S}^\theta]_{.i_1} \\ \vdots \\ [\hat{S}^\theta]_{.i_\alpha} \end{pmatrix} \tag{S5}$$

Also observe that, in the case of mass-action kinetics, equation (S3) can be simplified to [13]:

$$\boxed{\frac{\partial}{\partial t}\begin{pmatrix} \hat{p}^\theta \\ [\hat{S}^\theta]_{.i} \end{pmatrix} = \begin{pmatrix} \hat{A}^\theta & 0 \\ \hat{B}_i & \hat{A}^\theta \end{pmatrix}\begin{pmatrix} \hat{p}^\theta \\ [\hat{S}^\theta]_{.i} \end{pmatrix} \quad \forall i \in [\![1, M]\!]} \tag{S6}$$

where $\hat{A}^\theta \in \mathcal{M}_{N_{\max}, N_{\max}}(\mathbb{R})$ and with $[\hat{S}_0^\theta]_{.i} = 0$ as before, and $\hat{B}_i \in \mathcal{M}_{N_{\max}, N_{\max}}(\mathbb{R})$.

*Proof.* In this case, $\hat{A}^\theta$ can be written as a linear combination of $M$ $\theta$-independent matrices: $\hat{A}^\theta = \sum_{i=1}^{M} \theta_i \hat{B}_i$. We now have:

$$\forall i \in [\![1, M]\!], \quad \frac{\partial [\hat{S}]_{.i}}{\partial t}(\cdot; t, \theta) = \hat{B}_i \hat{p}(\cdot; t, \theta) + \hat{A}^\theta \big[\hat{S}\big]_{.i}(\cdot; t, \theta).$$

□

### S7 Computation of the Fisher Information

The Deep Learning and the Finite State Projection methods give approximations for the mass function and the sensitivity of the likelihood. These can then be used to numerically compute the infinite sum given by equation (2).

#### S7.1 Deep Learning method

The Fisher Information can be numerically approximated using $N_{\max}$ terms in the summation as:

$$\hat{\mathcal{I}}_t^\theta = [\hat{S}_t^\theta]^\top \operatorname{Diag}\left(\frac{1}{\hat{p}_t^\theta}\right)\hat{S}_t^\theta \tag{S7}$$

where we assimilate $\hat{p}_t^\theta$ to its image $\big\{\hat{p}_1(t,\theta), \hat{p}_2(t,\theta), ..., \hat{p}_{N_{\max}}(t,\theta)\big\}$.

This expression should be put in perspective with the following definition commonly given in Information Geometry [14]:

$$\boxed{\mathcal{I}_t^\theta = \Big\langle \nabla_\theta p(\cdot;t,\theta), \frac{1}{p(\cdot;t,\theta)} \nabla_\theta p(\cdot;t,\theta) \Big\rangle_{L^2(\mathbb{N}^N)}} \tag{S8}$$

*Proof.* If the state-space of $X_t$ is of cardinality $N_{\max}$, we can enumerate its elements as $\{x_t^1, x_t^2, ..., x_t^{N_{\max}}\}$. In this case, we have:

$$\begin{aligned}
\left[\mathcal{I}_t^\theta\right]_{ij} &= E_\theta\left[\left(\nabla_\theta \log p(X_t;t,\theta)\right)^\top \left(\nabla_\theta \log p(X_t;t,\theta)\right)\right]_{ij} \\
&= \sum_{\ell=1}^{N_{\max}} \Big[\left(\nabla_\theta \log p_\ell(t,\theta)\right)^\top \left(\nabla_\theta \log p_\ell(t,\theta)\right)\Big]_{ij} p_\ell(t,\theta) \\
&= \sum_{\ell=1}^{N_{\max}} \Big[\frac{\partial \log p_\ell}{\partial \theta_i}(t,\theta)\frac{\partial \log p_\ell}{\partial \theta_j}(t,\theta)\Big] p_\ell(t,\theta) \\
&= \sum_{\ell=1}^{N_{\max}} \Big(\frac{1}{p_\ell(t,\theta)}\frac{\partial p_\ell}{\partial \theta_i}(t,\theta)\Big)\Big(\frac{1}{p_\ell(t,\theta)}\frac{\partial p_\ell}{\partial \theta_j}(t,\theta)\Big) p_\ell(t,\theta) \\
&= \sum_{\ell=1}^{N_{\max}} \frac{1}{p_\ell(t,\theta)}\frac{\partial p_\ell}{\partial \theta_i}(t,\theta)\frac{\partial p_\ell}{\partial \theta_j}(t,\theta) \\
&= \sum_{\ell=1}^{N_{\max}} \frac{1}{p_\ell(t,\theta)} [S_t^\theta]_{\ell i} [S_t^\theta]_{\ell j} \\
&= \sum_{\ell=1}^{N_{\max}} [S_t^\theta]^\top_{i\ell} \Big[\mathrm{Diag}\Big(\frac{1}{p_\ell(t,\theta)}\Big) S_t^\theta\Big]_{\ell j} \\
&= \Big[[S_t^\theta]^\top \mathrm{Diag}\Big(\frac{1}{p_t^\theta}\Big) S_t^\theta\Big]_{ij}
\end{aligned}$$

□

##### S7.2 Finite State Projection method

For the Finite State Projection method, the solution of equation (S3) is used to compute the Fisher Information using formula (S7).

#### S8 Chemical Reaction Networks with piecewise-varying parameters

In the present study, we assume without loss of generality that one reaction is controlled. The uncontrolled reactions are defined in the same way as in section S3.

Let us define time points $\{t_l\}_{0 \le l \le L}$ such that $\{(t_{l-1}, t_l]\}_{1 \le l \le L}$ partitions $(0, t_f]$. The propensity $\lambda_M : \mathbb{N}^N \times \mathbb{R}_+^L \to \mathbb{R}_+$ of the controlled reaction depends on a non-negative, piecewise-constant control process whose value over the $L$ intervals is given by $\xi \in \mathbb{R}_+^L$:

$$\lambda_M(x_t;\xi) = \sum_{l=1}^{L} \tilde{\tilde{\lambda}}(x_t;\xi_l) \mathbb{1}_{(t_{l-1},t_l]}(t)$$

where: $\tilde{\tilde{\lambda}} : \mathbb{N}^N \times \mathbb{R}_+ \to \mathbb{R}_+$.

##### S8.1 Sampling with piecewise-varying parameters

The restarted Stochastic Simulation Algorithm uses the Markov property to simulate the trajectories sequentially over each time interval where parameters are constant.

**Algorithm S2** Restarted Stochastic Simulation Algorithm

**Input:** $x_0$, $s$, $\lambda$, $\theta$, $\{t_l\}$, $\xi$

1: $X \leftarrow x_0$, $t \leftarrow 0$

2: **for** $l \in [\![1, L]\!]$ **do**

3: $\quad \lambda_0 \leftarrow \sum_{k=1}^{M} \lambda_k(X; \theta, \xi)$

4: $\quad \Delta t \sim \mathcal{E}(\lambda_0)$

5: $\quad$ **if** $t + \Delta t > t_l$ **then**

6: $\quad\quad t \leftarrow t_l$

7: $\quad\quad X_l \leftarrow X$

8: $\quad$ **else**

9: $\quad\quad \Delta t_k \leftarrow \frac{\sum_{i=1}^{k} \lambda_i(X;\theta,\xi)}{\lambda_0}$ for $k \in [\![1, M]\!]$

10: $\quad\quad k^* \leftarrow \min\{k \in [\![1, M]\!] | \Delta t_k \geq \Delta t\}$

11: $\quad\quad t \leftarrow t + \Delta t$

12: $\quad\quad X \leftarrow X + s_{k^*}$

13: $\quad\quad$ Return to step (3)

14: $\quad$ **end if**

15: **end for**

**Output:** $\{X_{t_l}\}$

#### S8.2 Finite State Projection solution with piecewise-varying parameters

The restarted Finite State Projection algorithm solves the Chemical Master Equation sequentially over each time interval where parameters are constant.

**Algorithm S3** Restarted Finite State Projection

**Input:** $p_0$, $s$, $\lambda$, $\theta$, $\{t_l\}$, $\xi$

$l \leftarrow 0$

$\hat{p}_{t_l} \leftarrow p_0$, $\hat{S}_{t_l} \leftarrow 0$

**for** $l \in [\![1, L]\!]$ **do**

$\quad$ Solve equation (S5) for $\hat{p}_{t_l}^{\theta,\xi}$ and $\hat{S}_{t_l}^{\theta,\xi}$ over $[t_{l-1}, t_l]$ using $\hat{p}_{t_{l-1}}^{\theta,\xi}$ and $\hat{S}_{t_{l-1}}^{\theta,\xi}$ as initial conditions

**end for**

**Output:** $\{\hat{p}_{t_l}^{\theta,\xi}\}, \{\hat{S}_{t_l}^{\theta,\xi}\}$

### S9 Policy Search for Chemical Reaction Networks

We now explain how to use the mass function and sensitivity estimated by the Deep Learning and the Finite State Projection methods to perform policy search. Here, we directly place ourselves in the control framework presented in section S8.

#### S9.1 Performance indices for Chemical Reaction Networks

Recall that we consider the Optimal Control performance index $C_\xi^j$ defined in equation (3) by:

$$C_\xi^j = \sum_{l=1}^{L} w_l (E_{\theta,\xi}[X_{t_l}^j] - h_l)^2$$

Our aim is to find $\xi^*$, such that:

$$\xi^* = \arg\min_{\xi \in \mathcal{H}} C_\xi^j \tag{S9}$$

where $\mathcal{H} = \prod_{k=1}^{L} [a_k, b_k]$ with $a, b \in \mathbb{R}^L$ is a constraint set.

Note that our method is not limited to the performance index introduced in equation (3). Given a target mass function $p_c^{\theta,\xi}$ and a mass function $p^{\theta,\xi}$, an alternative is the Küllback-Leibler performance index $C_\xi^{\mathrm{KL},j}$ defined as:

$$C_\xi^{\mathrm{KL},j} = E_{p_c^{\theta,\xi}}\Big[\log\Big(\frac{p_c^{\theta,\xi}(X_t^j)}{p^{\theta,\xi}(X_t^j)}\Big)\Big] \tag{S10}$$

#### S9.2 Optimisation of performance indices for Reaction Networks

The constrained optimisation program (S9) is solved using Projected Gradient Descent, which projects optimisation parameters obtained by vanilla Gradient Descent onto the constraint interval $\mathcal{H}$.

**Algorithm S4** Projected Gradient Descent

**Input:** $n_{\text{iter}}$, $\varepsilon$, $\gamma$

$i \leftarrow 0$
$\xi \sim \mathcal{U}(\mathcal{H})$
**while** $i < n_{\text{iter}}$ and $\left\|\nabla_\xi C_\xi^J\right\|_2^2 > \varepsilon$ **do**
    $\xi \leftarrow \xi - \gamma \nabla_\xi C_\xi^J$
    **for** $k \in [\![1, L]\!]$ **do**
        **if** $\xi_k > b_k$ **then**
            $\xi_k \leftarrow b_k$
        **else if** $\xi_k < a_k$ **then**
            $\xi_k \leftarrow a_k$
        **end if**
    **end for**
    $i \leftarrow i + 1$
**end while**

Note that the optimisation algorithm is carried out using 3 hyperparameters: the maximal number of iterations $n_{\text{iter}}$, the threshold level $\varepsilon$ and the step size $\gamma$.

#### S9.3 Gradient of performance indices for Chemical Reaction Networks

The optimisation routine introduced above requires knowledge of $\nabla_\xi C_\xi^j$. The partial derivative of $C_\xi^j$ with respect to $\xi_i$ for $i \in [\![1, L]\!]$ is given by:

$$\frac{\partial C_\xi^j}{\partial \xi_i} = \sum_{l=1}^{L} 2w_l \frac{\partial E_{\theta,\xi}[X_{t_l}^j]}{\partial \xi_i}(E_{\theta,\xi}[X_{t_l}^j] - h_l) \tag{S11}$$

*Proof.*

$\forall i \in [\![1, L]\!]$,

$$\begin{aligned}\frac{\partial C_\xi^j}{\partial \xi_i} &= \frac{\partial}{\partial \xi_i}\Big(\sum_{l=1}^{L} w_l (E_{\theta,\xi}[X_{t_l}^j] - h_l)^2\Big) \\ &= \sum_{l=1}^{L} 2w_l \frac{\partial E_{\theta,\xi}[X_{t_l}^j]}{\partial \xi_i}(E_{\theta,\xi}[X_{t_l}^j] - h_l)\end{aligned}$$

□

The infinite sums involved in the expectations above can be numerically approximated using $N_{\max} \in \mathbb{N}^*$ terms, the partial derivative $\partial C_\xi^j / \partial \xi_i$ of the performance index being approximated as:

$$\frac{\widehat{\partial C_\xi^j}}{\partial \xi_i} = \sum_{l=1}^{L} 2w_l \Big( \sum_{k=1}^{N_{\max}} k[\hat{S}_{t_l}^{\theta,\xi}]_{k,M-1+i} \Big) ( \sum_{k=1}^{N_{\max}} k \ \hat{p}(k; t_l, \theta, \xi) - h_l)$$

*Proof.*

$$\begin{aligned}
\frac{\widehat{\partial C_\xi^j}}{\partial \xi_i} &= \sum_{l=1}^{L} 2w_l \frac{\partial}{\partial \xi_i} \Big( \sum_{k=1}^{N_{\max}} k \ \hat{p}(k; t_l, \theta, \xi) \Big) \Big( \sum_{k=1}^{N_{\max}} k \ \hat{p}(k; t_l, \theta, \xi) - h_l \Big) \\
&= \sum_{l=1}^{L} 2w_l \Big( \sum_{k=1}^{N_{\max}} k \ \frac{\partial \hat{p}(k; t_l, \theta, \xi)}{\partial \xi_i} \Big) \Big( \sum_{k=1}^{N_{\max}} k \ \hat{p}(k; t_l, \theta, \xi) - h_l \Big) \\
&= \sum_{l=1}^{L} 2w_l \Big( \sum_{k=1}^{N_{\max}} k[\hat{S}_{t_l}^{\theta,\xi}]_{k,M-1+i} \Big) \Big( \sum_{k=1}^{N_{\max}} k \ \hat{p}(k; t_l, \theta, \xi) - h_l \Big)
\end{aligned}$$

□

### S10 Formulas for the Fisher Information of selected uncontrolled Chemical Reaction Networks

In the study introducing Nessie, the Chemical Reaction Networks considered do not have known mass functions, and the accuracy of the Neural Network was hence directly tested against densities approximated using Monte Carlo estimation or the Finite State Projection. In contrast, in this section, we introduce three uncontrolled Chemical Reaction Networks with known mass functions: the Pure Production, Production and Degradation, and Explosive Chemical Reaction Networks. These are used to confirm that the mass functions estimated by the Mixture Density Network are accurate (see section S11).

The same Reaction Networks are then used to derive explicit formulas for sensitivities of the likelihood and Fisher Information diagonal elements. Similarly as before, we used these known formulas to test the accuracy of our method (see sections 3 and S11).

#### S10.1 Probability mass functions

##### S10.1.1 Production and Degradation Chemical Reaction Network

Recall that the Production and Degradation Chemical Reaction Network is defined by the following reaction graph:

$$\emptyset \underset{\theta_2}{\overset{\theta_1}{\rightleftharpoons}} S$$

Its dynamics are considered starting from the initial condition $x_0 = 0$, which means $P(X_0 = 0) = 1$.

$X_t$ then follows a Poisson distribution $\mathcal{P}(\lambda_t^\theta)$, where $\lambda_t^\theta = \frac{\theta_1}{\theta_2}(1 - e^{-\theta_2 t})$ [13]. This means that the probability of having $x$ elements of species $S$ at time $t$ is given by:

$$\boxed{p(x; t, \theta) = \frac{(\lambda_t^\theta)^x e^{-\lambda_t^\theta}}{x!}} \tag{S12}$$

##### S10.1.2 Pure Production Chemical Reaction Network

The Pure Production Chemical Reaction Network is defined by the following reaction graph:

$$\emptyset \xrightarrow{\theta} S$$

Its dynamics are considered starting from the initial condition $x_0 = 0$, which means $P(X_0 = 0) = 1$.

$X_t$ then follows a Poisson distribution $\mathcal{P}(\lambda_t^\theta)$, where $\lambda_t^\theta = \theta t$ [15]. This means that the probability of having $x$ elements of species $S$ at time $t$ is given by:

$$\boxed{p(x;t,\theta) = \frac{(\lambda_t^\theta)^x e^{-\lambda_t^\theta}}{x!}} \tag{S13}$$

##### S10.1.3 Explosive Production Chemical Reaction Network

The Explosive Production Chemical Reaction Network is defined by the following reaction graph:

$$S \xrightarrow{\theta} 2S$$

Its dynamics are considered starting from the initial condition $x_0 \in \mathbb{N}^*$, which means $P(X_0 = x_0) = 1$.

$X_t$ then follows a $x_0$-shifted Negative Binomial distribution $\mathcal{NB}(r, q_t^\theta)$, where $r = x_0$ and $q_t^\theta = e^{-t\theta}$ [16]. For $r \in \mathbb{N}^*$ and $q \in (0,1]$, we write $\mathcal{NB}(r,q)$ as $p_{\mathrm{NB}}(\cdot; r, q)$, and choose to interpret $r$ as the number of successes before the experiment is stopped, and to $q$ as the probability of success for each trial. It is equivalent to see $r$ as the number of failures before the experiment is stopped, and $q$ as the probability of failure for each trial. Under the same convention, the probability mass function of a Negative Binomial distribution is given by:

$$\forall k \in \mathbb{N}, \quad p_{\mathrm{NB}}(k; r, q) = \binom{k+r-1}{r-1}(1-q)^k q^r$$

The probability of having $x$ elements of species $S$ at time $t$ is then given by:

$$p(x;t,\theta) = p_{\mathrm{NB}}(x-r; r, q_t^\theta)\mathbb{1}_{x \geq r}$$

$$\boxed{p(x;t,\theta) = \binom{x-1}{r-1}\left(1-q_t^\theta\right)^{x-r}\left(q_t^\theta\right)^r \mathbb{1}_{x\geq r}} \tag{S14}$$

#### S10.2 Sensitivities of the likelihood

##### S10.2.1 Production and Degradation Chemical Reaction Network

The sensitivities of the likelihood of the Production and Degradation Chemical Reaction Network with respect to $\theta_1$ and $\theta_2$ are given by:

$$\boxed{\frac{\partial p}{\partial \theta_1}(x;t,\theta) = \frac{\lambda_t^\theta}{\theta_1}\big[p(x-1;t,\theta) - p(x;t,\theta)\big]} \tag{S15}$$

$$\boxed{\frac{\partial p}{\partial \theta_2}(x;t,\theta) = \Big(-\frac{\lambda_t^\theta}{\theta_2} + \frac{\theta_1}{\theta_2}te^{-\theta_2 t}\Big)\big[p(x-1;t,\theta) - p(x;t,\theta)\big]} \tag{S16}$$

*Proof.*

$\forall i \in \{1,2\}$,

$$\begin{aligned}
\frac{\partial p}{\partial \theta_i}(x;t,\theta) &= \frac{\partial \lambda_t^\theta}{\partial \theta_i}\frac{\partial p}{\partial \lambda_t^\theta}(x;t,\theta), \text{ by the chain rule}\\
&= \frac{\partial \lambda_t^\theta}{\partial \theta_i}\frac{\partial}{\partial \lambda_t^\theta}\left(\frac{(\lambda_t^\theta)^x e^{-\lambda_t^\theta}}{x!}\right) \text{ as } X_t \text{ follows a Poisson distribution of parameter } \lambda_t^\theta\\
&= \frac{\partial \lambda_t^\theta}{\partial \theta_i}\left(\frac{(\lambda_t^\theta)^{x-1} e^{-\lambda_t^\theta}}{(x-1)!} - \frac{(\lambda_t^\theta)^x e^{-\lambda_t^\theta}}{x!}\right)\\
&= \frac{\partial \lambda_t^\theta}{\partial \theta_i}\big[p(x-1;t,\theta) - p(x;t,\theta)\big]
\end{aligned}$$

This means:

$$\begin{aligned}\frac{\partial \lambda_t^\theta}{\partial \theta_1} &= \frac{1-e^{-\theta_2 t}}{\theta_2}\\ &= \frac{\lambda_t^\theta}{\theta_1}\end{aligned}$$

$$\begin{aligned}\frac{\partial \lambda_t^\theta}{\partial \theta_2} &= -\frac{\theta_1}{\theta_2^2}(1-e^{-\theta_2 t}) + \frac{\theta_1}{\theta_2} t e^{-\theta_2 t}\\ &= -\frac{\lambda_t^\theta}{\theta_2} + \frac{\theta_1}{\theta_2} t e^{-\theta_2 t}\end{aligned}$$

□

##### S10.2.2 Pure Production Chemical Reaction Network

The sensitivity of the likelihood of the Pure Production Chemical Reaction Network with respect to $\theta$ is given by:

$$\boxed{\frac{\partial p}{\partial \theta}(x;t,\theta) = t\big[p(x-1;t,\theta) - p(x;t,\theta)\big]} \tag{S17}$$

*Proof.*

Observe that:

$$\frac{\partial \lambda_t^\theta}{\partial \theta} = t$$

As in the previous proof, we then have:

$$\frac{\partial p}{\partial \theta}(x;t,\theta) = t\big[p(x-1;t,\theta) - p(x;t,\theta)\big]$$

□

##### S10.2.3 Explosive Production Chemical Reaction Network

The sensitivity of the likelihood of the Explosive Production Chemical Reaction Network with respect to $\theta$ is given by:

$$\boxed{\frac{\partial p}{\partial \theta}(x;t,\theta) = -t\ p(x;t,\theta)\Big[\frac{(r-x)q_t^\theta}{1-q_t^\theta} + r\Big]} \tag{S18}$$

*Proof.* First note that:

$$\frac{\partial q_t^\theta}{\partial \theta} = -te^{-\theta t} = -tq_t^\theta$$

This means that:

$$\begin{aligned}\frac{\partial p}{\partial \theta}(x;t,\theta) &= \frac{\partial q_t^\theta}{\partial \theta}\binom{x-1}{r-1}\Big[-(x-r)\big(1-q_t^\theta\big)^{x-r-1}\big(q_t^\theta\big)^r + \big(1-q_t^\theta\big)^{x-r} r \big(q_t^\theta\big)^{r-1}\Big]\mathbb{1}_{x\geq r}\\ &= \frac{\partial q_t^\theta}{\partial \theta}\binom{x-1}{r-1}\big(1-q_t^\theta\big)^{x-r}\big(q_t^\theta\big)^r\Big(\frac{r-x}{1-q_t^\theta} + \frac{r}{q_t^\theta}\Big)\mathbb{1}_{x\geq r}\\ &= -t\ p(x;t,\theta)\Big(\frac{(r-x)q_t^\theta}{1-q_t^\theta} + r\Big)\end{aligned}$$

□

#### S10.3 Fisher Information

##### S10.3.1 Production and Degradation Chemical Reaction Network

The diagonal elements of the Fisher Information of the Production and Degradation Chemical Reaction Network is given by:

$$\boxed{[\mathcal{I}_t^\theta]_{11} = \frac{\lambda_t^\theta}{\theta_1^2}} \tag{S19}$$

$$\boxed{[\mathcal{I}_t^\theta]_{22} = \Big(-\frac{1}{\theta_2} + \frac{t}{e^{\theta_2 t} - 1}\Big)^2 \lambda_t^\theta} \tag{S20}$$

*Proof.*

$\forall i \in \{1, 2\}$,

$$\begin{aligned}
[\mathcal{I}_t^\theta]_{ii} &= V_\theta\Big[\frac{\partial \log p}{\partial \theta_i}(X_t; t, \theta)\Big] \\
&= V_\theta\Big[\frac{1}{p(X_t; t, \theta)}\frac{\partial p}{\partial \theta_i}(X_t; t, \theta)\Big] \\
&= V_\theta\Big[\frac{\partial \lambda_t^\theta}{\partial \theta_i}\Big(\frac{X_t}{\lambda_t^\theta} - 1\Big)\Big]
\end{aligned}$$

$$\begin{aligned}
[\mathcal{I}_t^\theta]_{11} &= V_\theta\Big[\frac{\partial \lambda_t^\theta}{\partial \theta_1}\Big(\frac{X_t}{\lambda_t^\theta} - 1\Big)\Big] \\
&= V_\theta\Big[\frac{\lambda_t^\theta}{\theta_1}\Big(\frac{X_t}{\lambda_t^\theta} - 1\Big)\Big] \\
&= \frac{V_\theta[X_t]}{\theta_1^2} \\
&= \frac{\lambda_t^\theta}{\theta_1^2} \text{ as } X_t \text{ follows a Poisson distribution of parameter } \lambda_t^\theta
\end{aligned}$$

$$\begin{aligned}
[\mathcal{I}_t^\theta]_{22} &= V_\theta\Big[\frac{\partial \lambda_t^\theta}{\partial \theta_2}\Big(\frac{X_t}{\lambda_t^\theta} - 1\Big)\Big] \\
&= V_\theta\Big[\Big(-\frac{\lambda_t^\theta}{\theta_2} + \frac{\theta_1}{\theta_2} t e^{-\theta_2 t}\Big)\Big(\frac{X_t}{\lambda_t^\theta} - 1\Big)\Big] \\
&= \Big(-\frac{\lambda_t^\theta}{\theta_2} + \frac{\theta_1}{\theta_2} t e^{-\theta_2 t}\Big)^2 \frac{V_\theta[X_t]}{(\lambda_t^\theta)^2} \\
&= \frac{\Big[\Big(-\frac{1}{\theta_2} + \frac{t e^{-\theta_2 t}}{1 - e^{-\theta_2 t}}\Big)\lambda_t^\theta\Big]^2}{\lambda_t^\theta} \\
&= \Big(-\frac{1}{\theta_2} + \frac{t}{e^{\theta_2 t} - 1}\Big)^2 \lambda_t^\theta
\end{aligned}$$

□

##### S10.3.2 Pure Production Chemical Reaction Network

The Fisher Information of the Pure Production Chemical Reaction Network is given by:

$$\boxed{\mathcal{I}_t^\theta = \frac{t}{\theta}} \tag{S21}$$

*Proof.* As in the previous proof:

$$\begin{aligned}
\mathcal{I}_t^\theta &= V_\theta\Big[\frac{\partial \lambda_t^\theta}{\partial \theta}\Big(\frac{X_t}{\lambda_t^\theta} - 1\Big)\Big] \\
&= V_\theta\Big[t\Big(\frac{X_t}{\lambda_t^\theta} - 1\Big)\Big] \\
&= t^2 \frac{V_\theta[X_t]}{(\lambda_t^\theta)^2}
\end{aligned}$$

$$= \frac{t^2}{\lambda_t^\theta}$$
$$= \frac{t}{\theta}$$

□

##### S10.3.3 Explosive Production Chemical Reaction Network

The Fisher Information of the Explosive Production Chemical Reaction Network is given by:

$$\boxed{\mathcal{I}_t^\theta = \frac{rt^2}{1 - q_t^\theta}} \tag{S22}$$

*Proof.*

$$\mathcal{I}_t^\theta = V_\theta\Big[\frac{1}{p(X_t; t, \theta)}(-t)\ p(X_t; t, \theta)\Big(\frac{(r - X_t)q_t^\theta}{1 - q_t^\theta} + r\Big)\Big]$$
$$= V_\theta\Big[-t\Big(\frac{(r - X_t)q_t^\theta}{1 - q_t^\theta} + r\Big)\Big]$$
$$= \Big(\frac{tq_t^\theta}{1 - q_t^\theta}\Big)^2 V_\theta[X_t]$$
$$= \Big(\frac{tq_t^\theta}{1 - q_t^\theta}\Big)^2 \frac{r\big(1 - q_t^\theta\big)}{\big(q_t^\theta\big)^2}$$
$$= \frac{rt^2}{1 - q_t^\theta}$$

□

### S11 Additional numerical results for the computation of the Fisher Information

#### S11.1 Comparison of the time performance of the two methods

| | Pure Production | Production and Degradation | Explosive Production | Bursting Gene |
|---|---|---|---|---|
| Simulation time (SSA) for 4400 samples* ($h : min$) | 00 : 34 | 1 : 06 | 00 : 20 | 1 : 07 |
| Average training time** ($min : s$) | 4 : 12 | 5 : 04 | 16 : 50 | 13 : 34 |
| Average FSP runtime for the corresponding value of $C_r$*** ($s$) | 0.013 | 0.539 | 0.048 | 116 |

Table S2: Wall-clock computing time associated to the Deep Learning and Finite State Projection methods. * Each parameter configuration $\theta$ generates 4 samples, each of which corresponds to a sampling time $t_\ell$. Here, 1100 parameter configurations were generated, leading to the reported figure of 4400 samples. Simulations are run in parallel on 4 CPUs. ** Average over the training of 3 models. *** Average over 100 runs, for the value of $\theta_1$ used in the figures.

#### S11.2 Production and Degradation Chemical Reaction Network

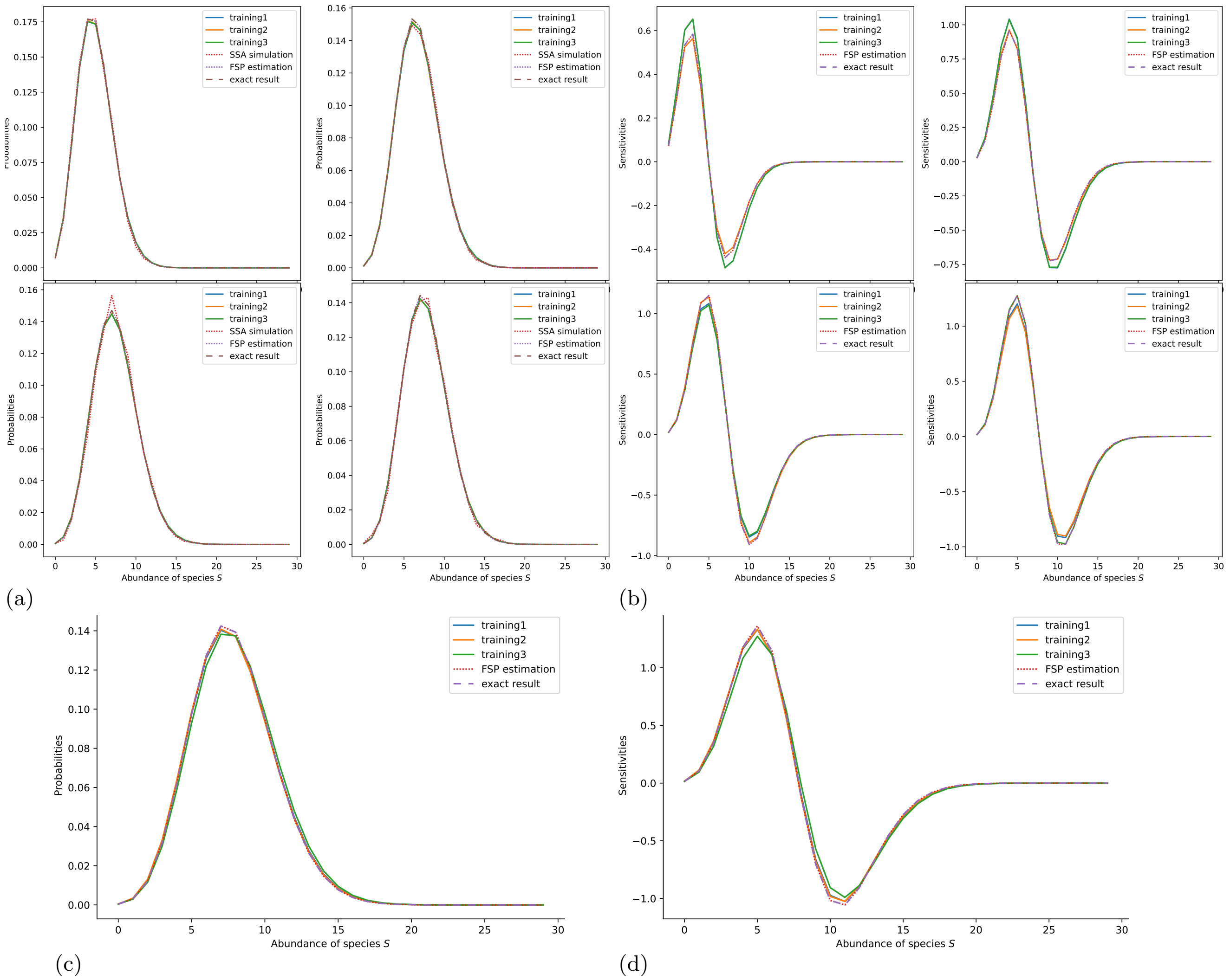

Figure S4: Probability mass function in figures S4a and S4c, and sensitivity of the likelihood with respect to $\theta_2$ in figures S4b and S4d for the Production and Degradation Chemical Reaction Network. Parameters: $\theta_1 = 1.5665, \theta_2 = 0.1997$. For figures S4a and S4b, from left to right, top to bottom, plots correspond to times $5, 10, 15, 20$. For figures S4c and S4d, time $t = 30$ is outside of the training range.

#### S11.3 Bursting Gene Chemical Reaction Network

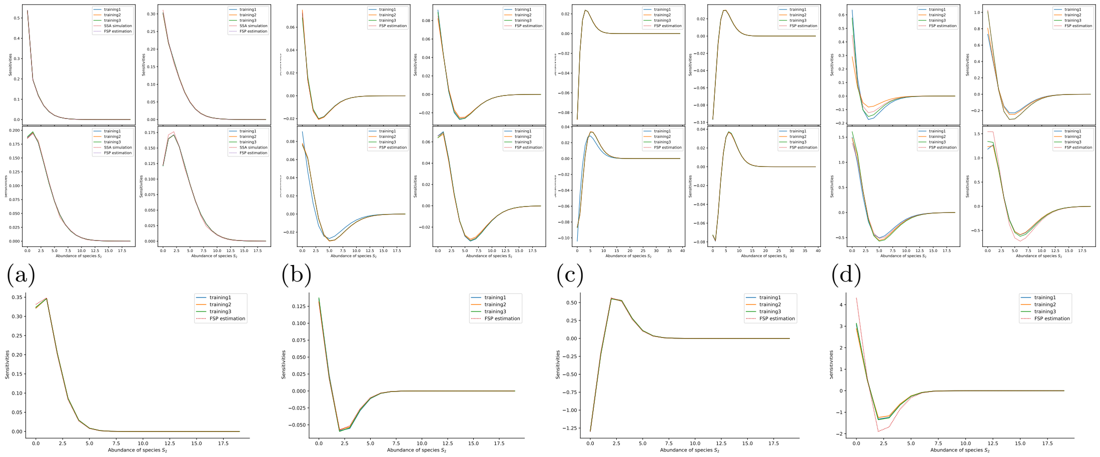

(a) (b) (c) (d)

(e) (f) (g) (h)

Figure S5: Probability mass function in figures S5a and S5e for the Bursting Gene Chemical Reaction Network. Sensitivity of the likelihood with respect to $\theta_2$ in figures S5b and S5f, with respect to $\theta_3$ in figures S5c and S5g, and with respect to $\theta_4$ in figures S5d and S5h for the Bursting Gene Chemical Reaction Network. Parameters: $\theta_1 = 0.6409, \theta_2 = 2.0359, \theta_3 = 0.2688, \theta_4 = 0.0368$. For figures S5a, S5b, S5c and S5d, from left to right, top to bottom, plots correspond to times $5, 10, 15, 20$. For figures S5e, S5f, S5g and S5h, time $t = 30$ is outside of the training range.

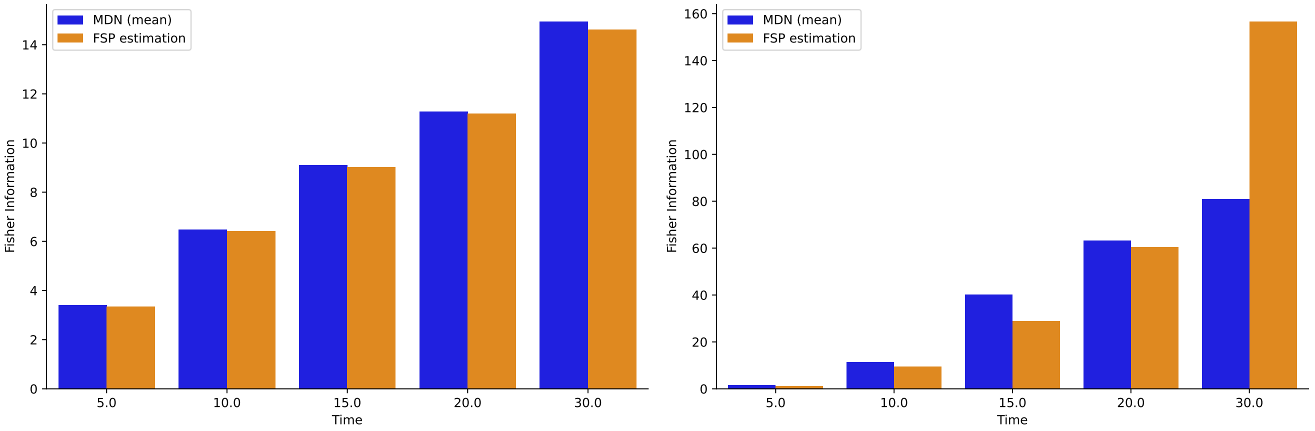

(a) (b)

Figure S6: Element $\left[\mathcal{I}_t^\theta\right]_{33}$ in figure S6a and $\left[\mathcal{I}_t^\theta\right]_{44}$ in figure S6b of the Fisher Information for the Bursting Gene Chemical Reaction Network as a function of time. Parameters: $\theta_1 = 0.6409, \theta_2 = 2.0359, \theta_3 = 0.2688, \theta_4 = 0.0368$. Time $t = 30$ is outside of the training range.

#### S11.4 Toggle Switch Chemical Reaction Network

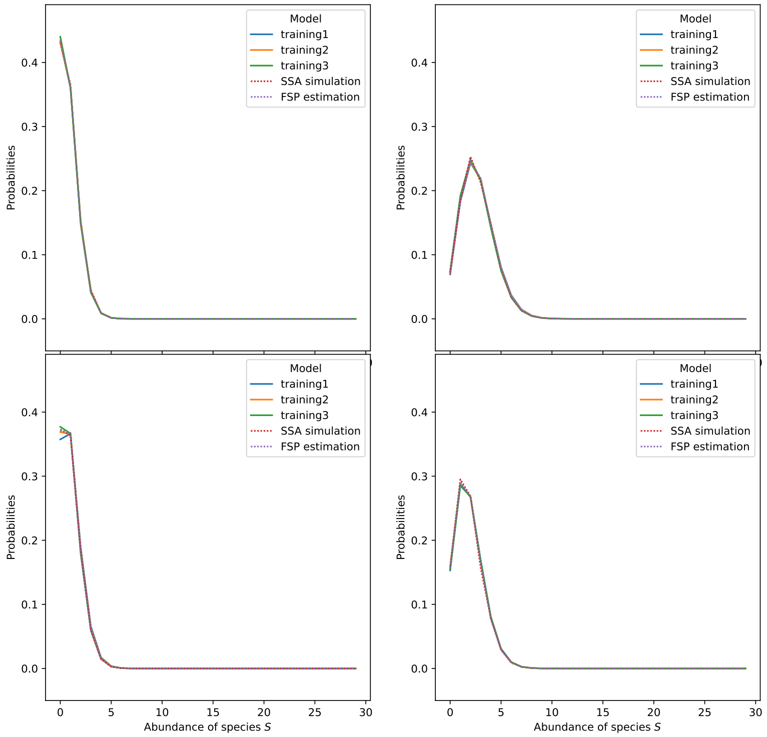

Figure S7: Probability mass function for the Toggle Switch Chemical Reaction Network. Parameters: $\theta_1 = 0.7455, \theta_2 = 0.3351, \theta_3 = 0.0078, \theta_4 = 0.4656, \theta_5 = 0.0193, \theta_6 = 0.2696, \theta_7 = 2.5266, \theta_8 = 0.4108, \theta_9 = 0.6880, \xi_1 = 0.9276, \xi_2 = 0.2132, \xi_3 = 0.8062, \xi_4 = 0.3897$. From left to right, top to bottom, plots correspond to times $5, 10, 15, 20$.

#### S11.5 Pure Production Chemical Reaction Network

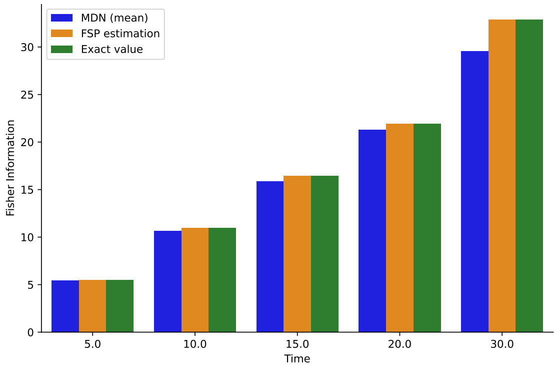

Figure S8: Fisher Information as a function of time for the Pure Production Chemical Reaction Network. Parameter: $\theta = 0.9120$. Time $t = 30$ is outside of the training range.

#### S11.6 Explosive Production Chemical Reaction Network

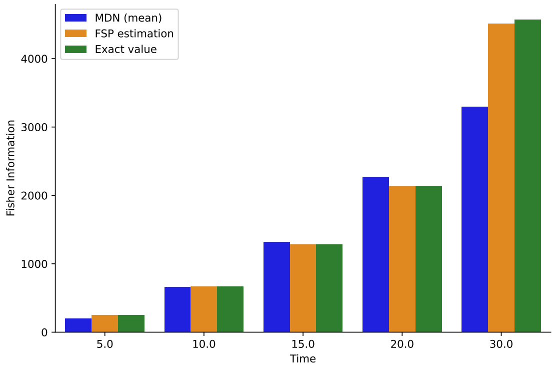

Figure S9: Fisher Information as a function of time for the Explosive Production Chemical Reaction Network. Parameter: $\theta = 0.1384$. Time $t = 30$ is outside of the training range.

#### S11.7 Values used for the numerical results

| | Pure Production | Production and Degradation | Explosive Production | Bursting Gene | Toggle Switch |
|---|---|---|---|---|---|
| $x_0$ | 0 | 0 | 5 | $(0, 0)$ | $(0, 0)$ |
| $\theta$ | $\theta \in (0, 2]$ | $\theta_1 \in (0, 2], \theta_2 \in (0, 1]$ | $\theta \in (0, 0.2]$ | $\theta_1 \in (0, 1], \theta_2 \in (0, 3], \theta_3 \in (0, 5], \theta_4 \in (0, 0.05]$ | $(\theta_1, \theta_2, \theta_3, \theta_4, \theta_5, \theta_6) \in (0, 1]^6, \theta_7 \in (0, 2], \theta_9 \in (0, 3], \theta_9 \in (0, 1]$ |
| $\xi$ | - | - | - | - | $\xi \in (0, 1]$ |
| $j$ | 1 | 1 | 1 | 2 | 2 |
| $t_\ell$ | $t_1 = 5, t_2 = 10, t_3 = 15, t_4 = 20$ | | | | |

Table S3: Parameters used to specify the Chemical Reaction Networks on which to train the Mixture Density Networks.

| Hyperparameter | Pure Production | Production and Degradation | Explosive Production | Bursting Gene | Toggle Switch |
|---|---|---|---|---|---|
| $n_{\text{sampled}}$ | 4400 | 4400 | 4400 | 8800 | 42624 |
| $n_{\text{sim}}$ | $10^4$ | $10^4$ | $10^4$ | $10^4$ | $10^4$ |
| $K$ | 4 | 4 | 4 | 4 | 4 |
| $N_{\text{hidden}}$ | 128 | 256 | 256 | 128 | 256 |
| $n_{\text{batches}}$ | 32 | 32 | 64 | 32 | 32 |
| $l_{r,0}$ | 0.005 | 0.005 | 0.005 | 0.005 | 0.001 |
| $n_{\text{epochs}}$ | 700 | 700 | 500 | 700 | 700 |

Table S4: Hyperparameters chosen to train the Mixture Density Network on the Chemical Reaction Networks presented.

| Hyperparameter | Pure Production | Production and Degradation | Explosive Production | Bursting Gene | Toggle Switch |
|---|---|---|---|---|---|
| $N_{\max}$ (MDN) | 200 | 200 | 1000 | 200 | 200 |
| $C_r$ (FSP) | 200 | 200 | 1000 | 400 | 400 |
| *Corresponding* $N_{\max}$ *(FSP)* | 200 | 200 | 1000 | 80600 | 1325 |

Table S5: Hyperparameters chosen for the numerical computation of the infinite sum in equation (2).

Note that $N_{\max}$ could be made to vary with the value of the parameters considered.

### S12 Additional numerical results for the computation of the performance index and its gradient

#### S12.1 Comparison of the time performance of the two methods

| | Controlled Pure Production | Controlled Bursting Gene | Toggle Switch |
|---|---|---|---|
| Simulation time (SSA) for 4400 samples* ($h : min$) | 1 : 08 | 3 : 18 | 4 : 28 |
| Average training time** ($min : s$) | 9 : 22 | 29 : 42 | 49 : 25 |
| Average FSP runtime for the corresponding value of $C_r$*** ($s$) | 0.27 | 144 | 154 |

Table S6: Wall-clock computing time associated to the Deep Learning and Finite State Projection methods. * Each parameter configuration $\theta$ generates 4 samples, each of which corresponds to a sampling time $t_\ell$. Here, 1100 parameter configurations were generated, leading to the reported figure of 4400 samples. Simulations are run in parallel on 4 CPUs. ** Average over the training of 3 models.*** Average over 100 runs, for the value of $\theta_1$ used in the figures.

#### S12.2 Controlled Production and Degradation Chemical Reaction Network

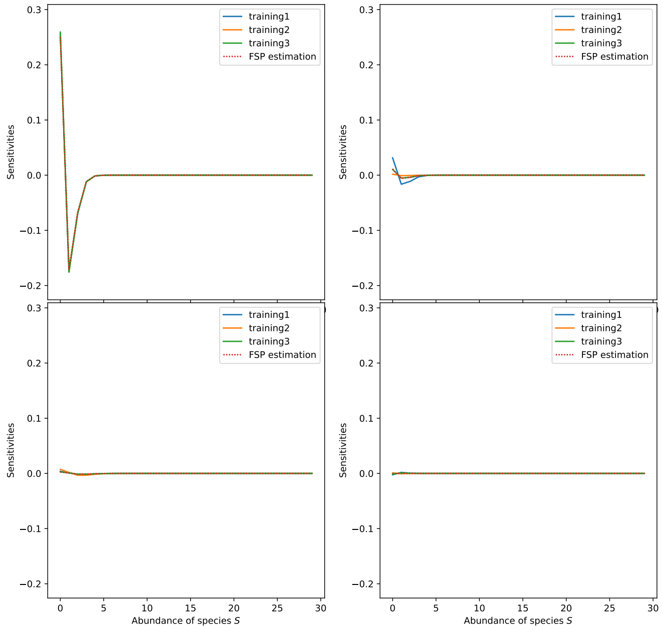

(a)

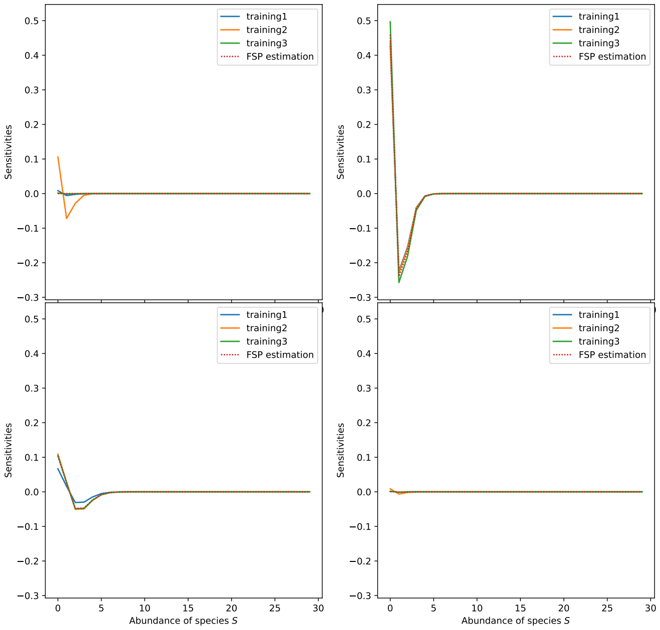

(b)

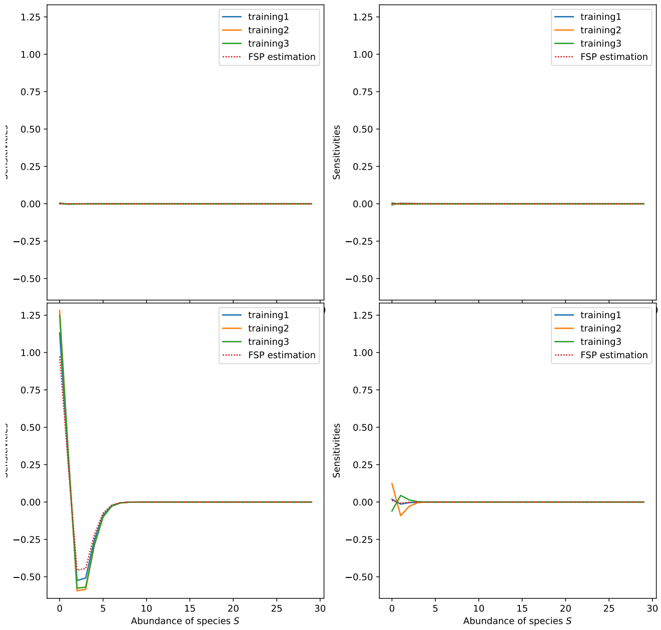

(c)

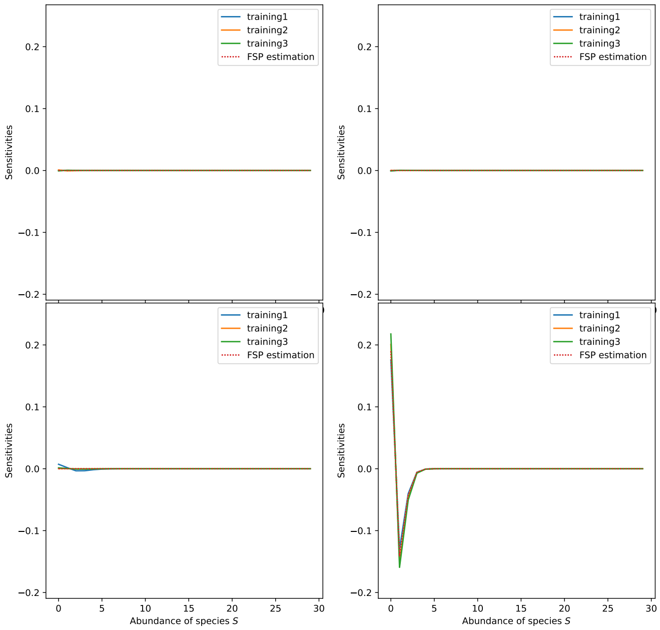

(d)

Figure S10: Sensitivity of the likelihood with respect to $\xi_1$ in figure S10a, $\xi_2$ in figure S10b, $\xi_3$ in figure S10c, and $\xi_4$ in figure S10d for the controlled Production and Degradation Chemical Reaction Network. For each figure, from left to right, top to bottom, plots correspond to times $5, 10, 15, 20$. Parameters: $\theta = 1.6869, \xi_1 = 1.8234, \xi_2 = 0.3082, \xi_3 = 0.1011, \xi_4 = 1.7771$.

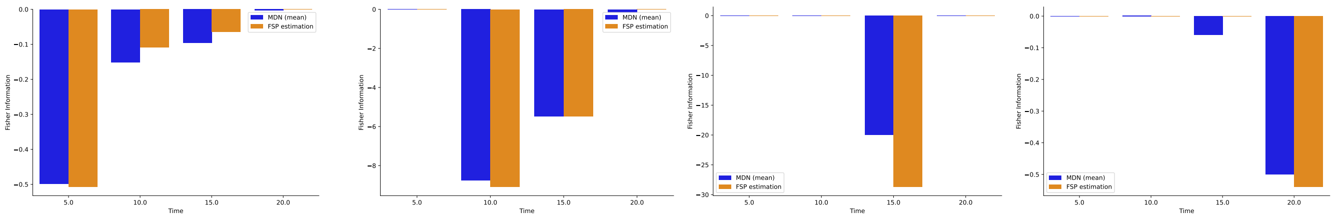

(a) (b) (c) (d)

Figure S11: Gradient of the expectation with respect to $\xi_1$ in figure S11a, $\xi_2$ in figure S11b, $\xi_3$ in figure S11c, and $\xi_4$ in figure S11d as a function of time for the controlled Production and Degradation Chemical Reaction Network. Parameters: $\theta = 1.6869, \xi_1 = 1.8234, \xi_2 = 0.3082, \xi_3 = 0.1011, \xi_4 = 1.7771$.

#### S12.3 Controlled Bursting Gene

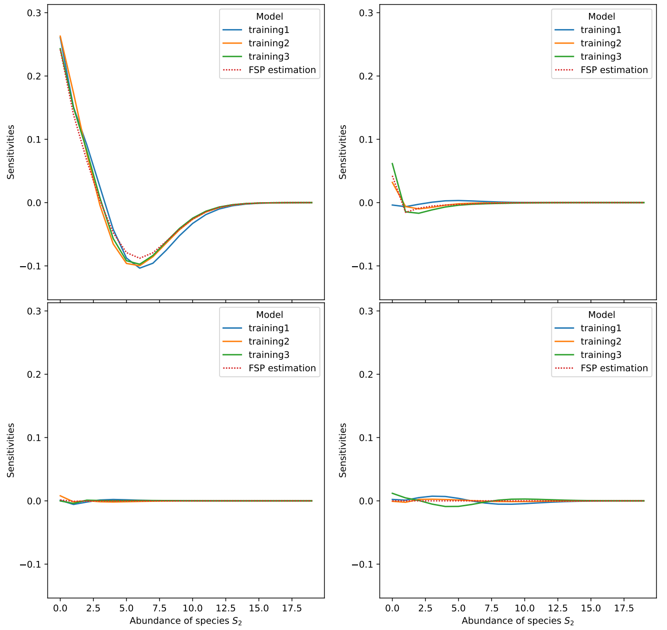

(a)

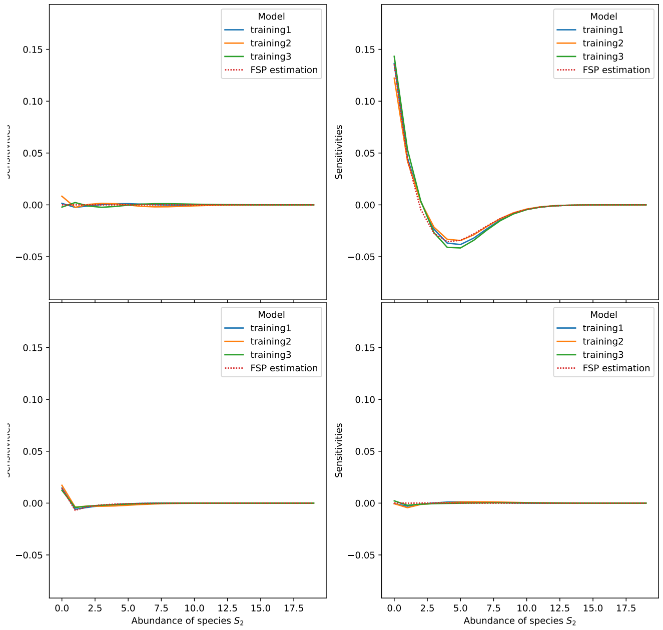

(b)

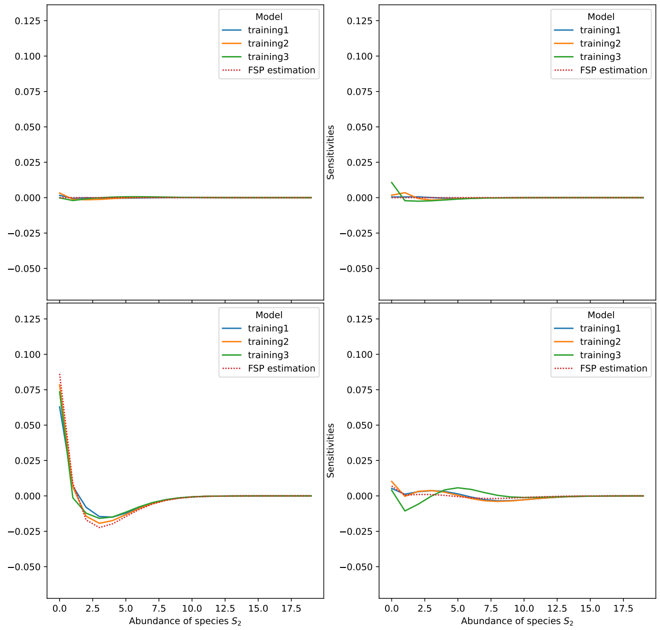

(c)

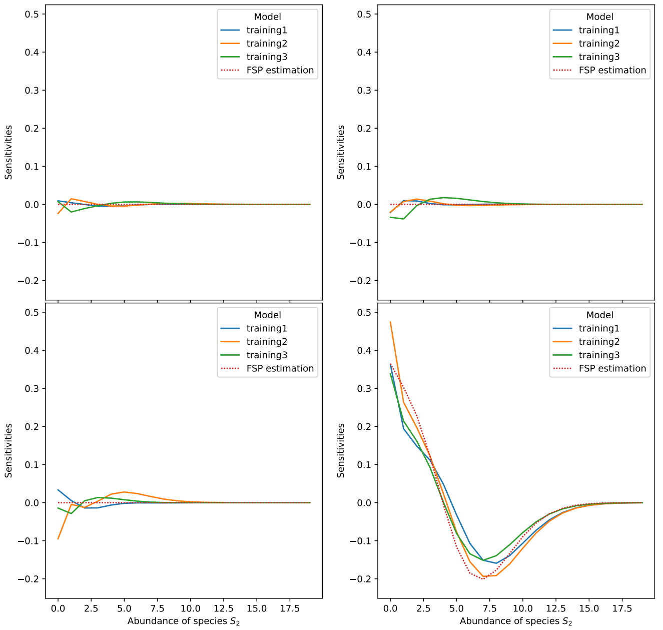

(d)

Figure S12: Sensitivity of the likelihood with respect to $\xi_1$ in figure S12a, $\xi_2$ in figure S12b, $\xi_3$ in figure S12c), and $\xi_4$ in figure S12d for the controlled Bursting Gene Chemical Reaction Network. For each figure, from left to right, top to bottom, plots correspond to times $5, 10, 15, 20$. Parameters: $\theta_1 = 0.4622, \theta_2 = 4.9699, \theta_3 = 0.7501, \xi_1 = 0.4677, \xi_2 = 1.3301, \xi_3 = 2.2814, \xi_4 = 0.0594$.

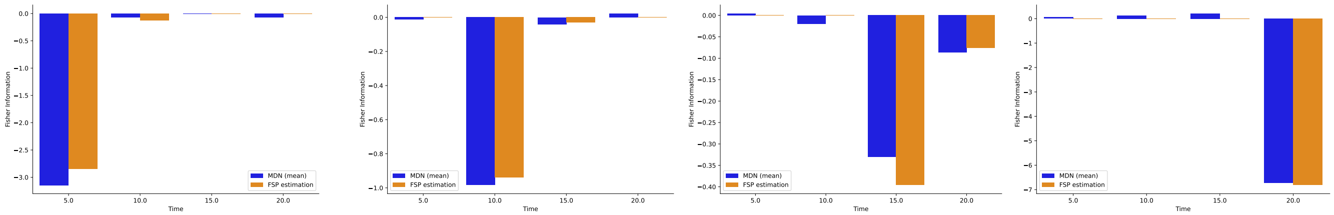

(a) (b) (c) (d)

Figure S13: Gradient of the expectation with respect to $\xi_1$ in S13a, $\xi_2$ in figure S13b, $\xi_3$ in figure S13c, and $\xi_4$ in figure S13d as a function of time for the controlled Bursting Gene Chemical Reaction Network. Parameters: $\theta_1 = 0.4622, \theta_2 = 4.9699, \theta_3 = 0.7501, \xi_1 = 0.4677, \xi_2 = 1.3301, \xi_3 = 2.2814, \xi_4 = 0.0594$.

#### S12.4 Toggle Switch Chemical Reaction Network

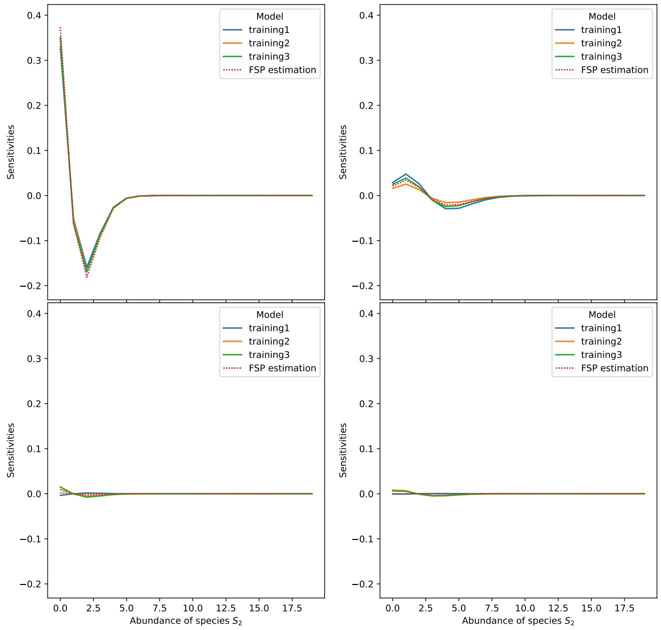

(a)

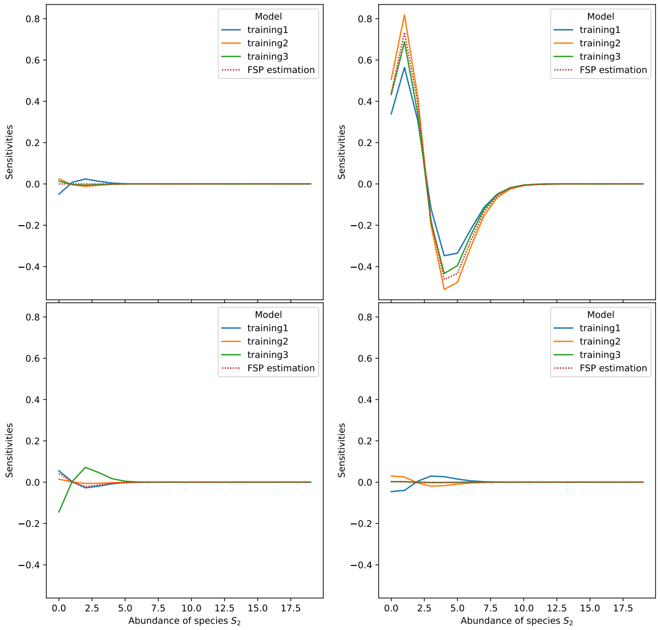

(b)

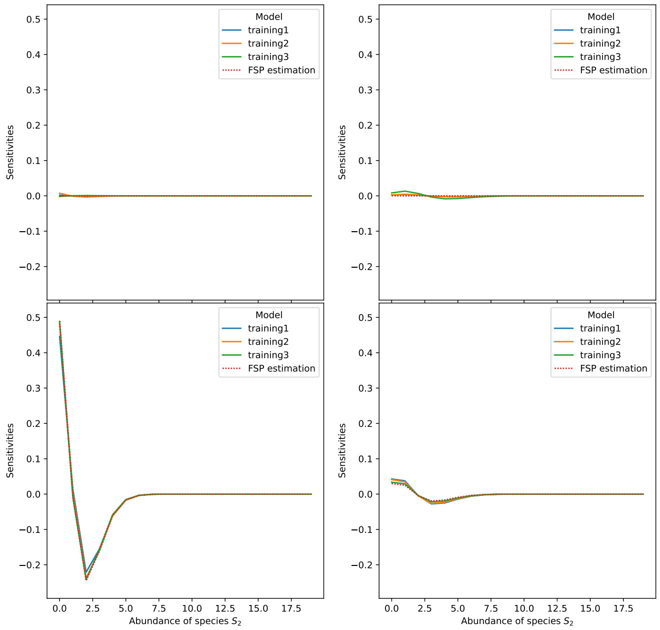

(c)

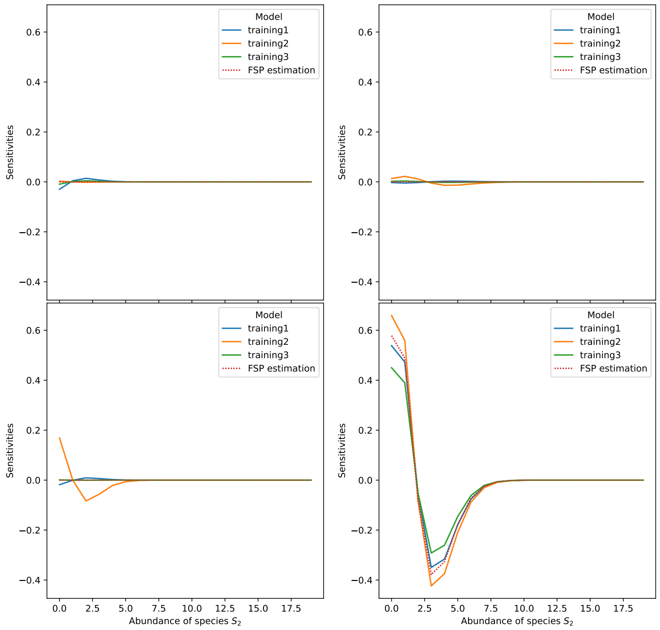

(d)

Figure S14: Sensitivity of the likelihood with respect to $\xi_1$ in figure S14a, $\xi_2$ in figure S14b, $\xi_3$ in figure S14c, and $\xi_4$ in figure S14d for the Toggle Switch Chemical Reaction Network. For each figure, from left to right, top to bottom, plots correspond to times $5, 10, 15, 20$. Parameters: $\theta_1 = 0.7455, \theta_2 = 0.3351, \theta_3 = 0.0078, \theta_4 = 0.4656, \theta_5 = 0.0193, \theta_6 = 0.2696, \theta_7 = 2.5266, \theta_8 = 0.4108, \theta_9 = 0.6880, \xi_1 = 0.9276, \xi_2 = 0.2132, \xi_3 = 0.8062, \xi_4 = 0.3897$.

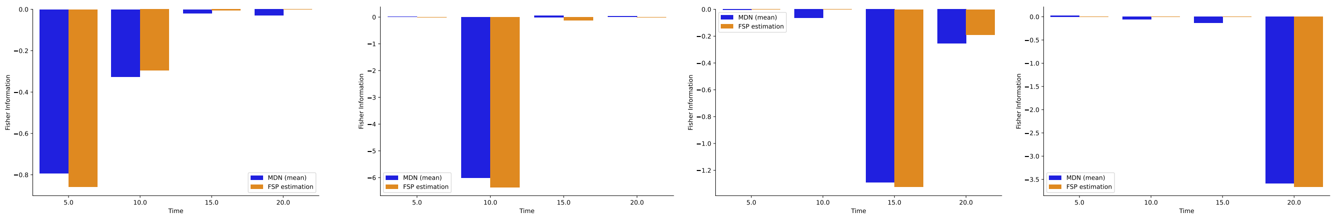

(a) (b) (c) (d)

Figure S15: Gradient of the expectation with respect to $\xi_1$ in figure S15a, $\xi_2$ in figure S15b, $\xi_3$ in figure S15c, and $\xi_4$ in figure S15d as a function of time for the Toggle Switch Chemical Reaction Network. Parameters: $\theta_1 = 0.7455, \theta_2 = 0.3351, \theta_3 = 0.0078, \theta_4 = 0.4656, \theta_5 = 0.0193, \theta_6 = 0.2696, \theta_7 = 2.5266, \theta_8 = 0.4108, \theta_9 = 0.6880, \xi_1 = 0.9276, \xi_2 = 0.2132, \xi_3 = 0.8062, \xi_4 = 0.3897$.

##### S12.5 Values used for the numerical results

| | Controlled Production and Degradation | Controlled Bursting Gene | Toggle Switch |
|---|---|---|---|
| $x_0$ | 0 | $(0,0)$ | $(0,0)$ |
| $\theta$ | $\theta \in (0,2]$ | $\theta_1 \in (0,2], \theta_2 \in (0,5], \theta_3 \in (0,2]$ | $(\theta_1,\theta_2,\theta_3,\theta_4,\theta_5,\theta_6) \in (0,1]^6, \theta_7 \in (0,2], \theta_8 \in (0,3], \theta_9 \in (0,1]$ |
| $\xi$ | $\xi \in (0,2]$ | $\xi \in (0,3]$ | $\xi \in (0,1]$ |
| $J$ | 1 | 2 | 2 |
| $t_\ell$ | $t_1 = 5, t_2 = 10, t_3 = 15, t_4 = 20$ | | |

Table S7: Parameters chosen to train the Mixture Density Network on the controlled Chemical Reaction Networks presented.

| Hyperparameter | Controlled Production and Degradation | Controlled Bursting Gene | Toggle Switch |
|---|---|---|---|
| $n_{\text{sampled}}$ | 9856 | 20776 | 42624 |
| $n_{\text{sim}}$ | $10^4$ | $10^4$ | $10^4$ |
| $K$ | 4 | 4 | 4 |
| $N_{\text{hidden}}$ | 256 | 128 | 256 |
| $n_{\text{batches}}$ | 32 | 32 | 32 |
| $l_{r,0}$ | 0.005 | 0.005 | 0.001 |
| $n_{\text{epochs}}$ | 700 | 700 | 700 |
| $N_{\text{max}}$ | 200 | | |

Table S8: Hyperparameters chosen to train Mixture Density Network on the controlled Chemical Reaction Networks presented.

### S13 Additional numerical results for the policy search

##### S13.1 Comparison of the performance of the two methods

| Controlled Production and Degradation | | | | | | | | |
|---|---|---|---|---|---|---|---|---|
| Control task | (9a) | (S16a) | (S16b) | (S16c) | (9b) | (9c) | (S16d) | (9d) |
| Effective number of iterations | 4046 | 269 | 473 | 1490 | 1793 | 18479 | 1866 | 1294 |
| Runtime (s) | 48.2 | 3.71 | 5.47 | 19.1 | 21.5 | 222 | 22.5 | 15.0 |
| Final loss (estimated by the MDN) | $1.02 \cdot 10^{-7}$ | $5.35 \cdot 10^{-9}$ | $3.63 \cdot 10^{-9}$ | $2.42 \cdot 10^{-10}$ | $6.45 \cdot 10^{-8}$ | $1.73 \cdot 10^{-7}$ | $5.66 \cdot 10^{-8}$ | $4.78 \cdot 10^{-8}$ |
| Final loss (estimated by the SSA*) | $7.86 \cdot 10^{-4}$ | $12.8 \cdot 10^{-3}$ | $3.56 \cdot 10^{-3}$ | $1.35 \cdot 10^{-2}$ | $6.07 \cdot 10^{-4}$ | $1.18 \cdot 10^{-2}$ | $8.67 \cdot 10^{-3}$ | $2.02 \cdot 10^{-2}$ |
| Estimated $\xi_1^*$ | 2.002 | 0.9970 | 0.6568 | 0.4449 | 2.003 | 0.6617 | 0.6623 | 0.6703 |
| Estimated $\xi_2^*$ | 1.999 | 0.9965 | 0.6611 | 0.5091 | 0.9968 | 0.9830 | 0.9902 | 1.9601 |
| Estimated $\xi_3^*$ | 2.012 | 1.000 | 0.6701 | 0.5080 | 0.9974 | 2.002 | 1.987 | 0.6429 |
| Estimated $\xi_4^*$ | 2.013 | 0.9975 | 0.6632 | 0.5020 | 0.6582 | 4.079 | 0.6561 | 2.007 |

Table S9: Results of the policy search for the controlled Production and Degradation Chemical Reaction Network using the Mixture Density Network. *Expectation calculated from $10^4$ samples obtained from the Stochastic Simulation Algorithm.

| Controlled Bursting Gene | | | | | | | | |
|---|---|---|---|---|---|---|---|---|
| Control task | (S17a) | (10a) | (S17b) | (S17c) | (10b) | (S17d) | (10c) | (10d) |
| Effective number of iterations | 26887 | 5428 | 916 | 50000 | 30000 | 22549 | 30000 | 30000 |
| Runtime (s) | 371 | 77.5 | 10.9 | 608 | 400 | 261 | 420 | 424 |
| Final loss (estimated by the MDN) | $2.30 \cdot 10^{-7}$ | $1.09 \cdot 10^{-8}$ | $2.23 \cdot 10^{-9}$ | $6.21 \cdot 10^{-3}$ | $8.63 \cdot 10^{-6}$ | $2.29 \cdot 10^{-7}$ | $4.26 \cdot 10^{-7}$ | $1.87 \cdot 10^{-3}$ |
| Final loss (estimated by the SSA*) | $1.65 \cdot 10^{-3}$ | $1.62 \cdot 10^{-3}$ | $1.78 \cdot 10^{-2}$ | $4.39 \cdot 10^{-2}$ | $4.72 \cdot 10^{-3}$ | $2.88 \cdot 10^{-2}$ | $5.72 \cdot 10^{-2}$ | $5.08 \cdot 10^{-2}$ |
| Estimated $\xi_1^*$ | 3.081 | 0.9766 | 0.2044 | 0.02172 | 3.194 | 3.035 | 0.02716 | 0.2122 |
| Estimated $\xi_2^*$ | 2.815 | 0.9837 | 0.3560 | 0.2250 | 0.8914 | 0.8260 | 0.4917 | 4.829 |
| Estimated $\xi_3^*$ | 2.772 | 1.006 | 0.3500 | 0.1917 | 0.9902 | 0.2139 | 1.201 | 0.1986 |
| Estimated $\xi_4^*$ | 3.034 | 0.9718 | 0.4030 | 0.2420 | 0.3000 | 0.1303 | 3.526 | 3.376 |

Table S10: Results of the policy search for the controlled Bursting Gene Chemical Reaction Network using the Mixture Density Network. *Expectation calculated from $10^4$ samples obtained from the Stochastic Simulation Algorithm.

| Toggle Switch | | | | | | | | |
|---|---|---|---|---|---|---|---|---|
| Control trask | (S18a) | (S18b) | (11a) | (S18c) | (11b) | (S18d) | (11c) | (11d) |
| Effective number of iterations | 5328 | 2292 | 30000 | 11022 | 281 | 170 | 1372 | 1413 |
| Runtime (s) | 61.9 | 41.4 | 420 | 127 | 3.29 | 2.81 | 16.2 | 17.5 |
| Final loss (estimated by the MDN) | $1.29 \cdot 10^{-7}$ | $2.00 \cdot 10^{-11}$ | $8.04 \cdot 10^{-9}$ | $3.52 \cdot 10^{-11}$ | $8.01 \cdot 10^{-10}$ | $4.18 \cdot 10^{-9}$ | $4.70 \cdot 10^{-8}$ | $4.33 \cdot 10^{-8}$ |
| Final loss (estimated by the SSA*) | $5.76 \cdot 10^{-2}$ | $4.27 \cdot 10^{-2}$ | $2.28 \cdot 10^{-2}$ | $3.17 \cdot 10^{-2}$ | $3.12 \cdot 10^{-2}$ | $3.92 \cdot 10^{-2}$ | $8.29 \cdot 10^{-2}$ | $4.97 \cdot 10^{-1}$ |
| Estimated $\xi_1^*$ | 1.066 | 0.7518 | 0.4727 | 0.2929 | 0.745 | 1.027 | 0.3101 | 0.2377 |
| Estimated $\xi_2^*$ | 1.030 | 0.7748 | 0.5224 | 0.4036 | 0.507 | 0.7489 | 0.5115 | 1.530 |
| Estimated $\xi_3^*$ | 1.076 | 0.7979 | 0.5286 | 0.3970 | 0.528 | 0.5063 | 0.8614 | 0.3645 |
| Estimated $\xi_4^*$ | 1.070 | 0.8119 | 0.5332 | 0.4058 | 0.393 | 0.3935 | 1.544 | 1.522 |

Table S11: Results of the policy search for the Toggle Switch Chemical Reaction Network using the Mixture Density Network. *Expectation calculated from $10^4$ samples obtained from the Stochastic Simulation Algorithm.

| CRN | Controlled Production and Degradation | Controlled Bursting Gene | Toggle Switch |
|---|---|---|---|
| Control task | (9a) | (S17a) | (S18a) |
| $n_{\text{iter}}$ | 20000 | 30000 | 50000 |
| $\gamma$ | 0.01 | 0.01 | 0.001 |
| $\varepsilon$ | $10^{-7}$ | $10^{-8}$ | $10^{-7}$ |
| Minimal loss | $7.86 \cdot 10^{-4}$ | $1.65 \cdot 10^{-3}$ | $5.76 \cdot 10^{-2}$ |
| $C_r$ | 100 | 100 | 100 |
| Effective number of iterations | 1822 | 5902 | 10287 |
| Runtime (s) | 1518 | 70456 | 50995 |
| Runtime FSP/Runtime MDN | 31.5 | 190 | 824 |
| Final loss (estimated by the FSP) | $7.81 \cdot 10^{-4}$ | $1.65 \cdot 10^{-3}$ | $5.75 \cdot 10^{-2}$ |
| Estimated $\xi_1^*$ | 2.002 | 2.886 | 0.9806 |
| Estimated $\xi_2^*$ | 2.042 | 3.025 | 1.161 |
| Estimated $\xi_3^*$ | 2.039 | 3.317 | 0.9809 |
| Estimated $\xi_4^*$ | 2.000 | 2.883 | 0.9848 |

Table S12: Results of the policy search using the Finite State Project method. $J$, $\mathcal{H}$, $h$, $n_{\text{iter}}$ are the same as in the equivalent experiment run with the Mixture Density Network.

#### S13.2 Controlled Production and Degradation Chemical Reaction Network

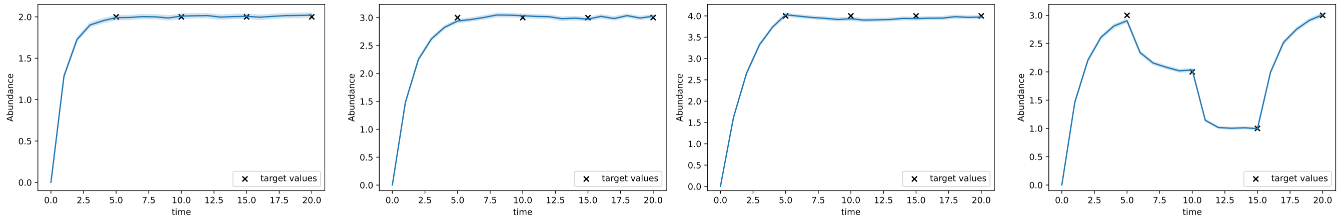

(a) (b) (c) (d)

Figure S16: Evolution of the expected abundance of $S$ for the controlled Production and Degradation Chemical Reaction Network using the $\xi^*$ obtained by policy search. Results, parameters, tasks and hyperparameters are detailed in tables S9, S13, S14 and S15.

#### S13.3 Controlled Bursting Gene Chemical Reaction Network

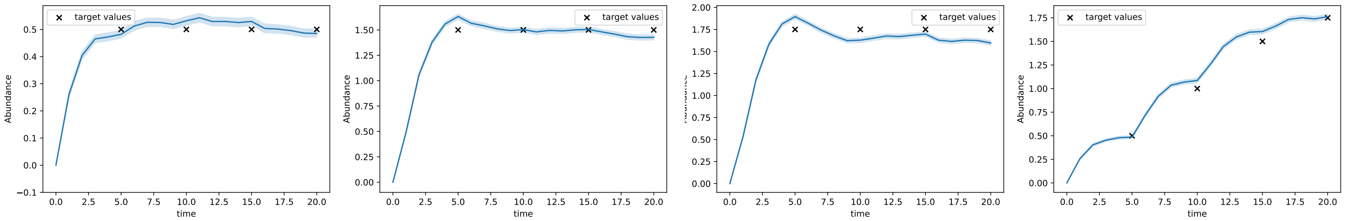

(a) (b) (c) (d)

Figure S17: Evolution of the expected abundance of $S_2$ for the controlled Bursting Gene Chemical Reaction Network using the $\xi^*$ obtained by policy search. Results, parameters, tasks and hyperparameters are detailed in tables S10, S13, S16 and S17.

#### S13.4 Toggle Switch Chemical Reaction Network

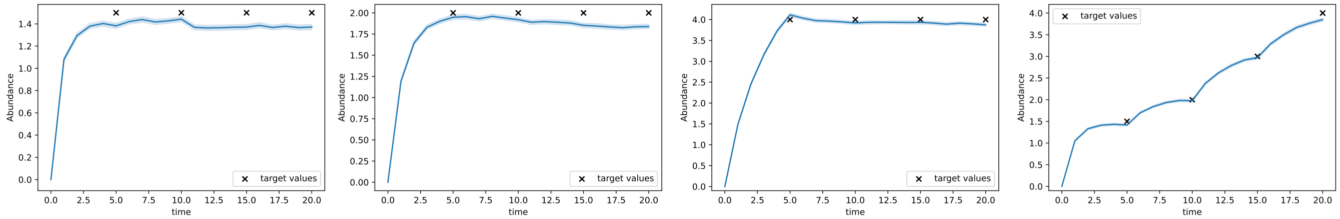

(a) (b) (c) (d)

Figure S18: Evolution of the expected abundance of $S_2$ for the Toggle Switch Chemical Reaction Network using the parameters $\xi^*$ obtained by policy search. Results, parameters, tasks and hyperparameters are detailed in tables S11, S13, S18 and S19.

#### S13.5 Values used for the numerical results

| | Controlled Production and Degradation | Controlled Bursting Gene | Toggle Switch |
|---|---|---|---|
| $x_0$ | 0 | $(0, 0)$ | $(0, 0)$ |
| $\theta$ | $\theta = 2$ | $\theta_1 = 1, \theta_2 = 2, \theta_3 = 1$ | $\theta_1 = 1, \theta_2 = 1, \theta_3 = 1, \theta_4 = 1, \theta_5 = 1, \theta_6 = 1, \theta_7 = 2.1, \theta_8 = 3, \theta_9 = 1$ |
| $t_\ell$ | $t_1 = 5, t_2 = 10, t_3 = 15, t_4 = 20$ | | |

Table S13: Parameters used to specify the controlled Chemical Reaction Networks on which the Optimal Control experiments are performed. Recall that the initial state is not an input of the Mixture Density Network.

| Controlled Production and Degradation | | | | | | | | |
|---|---|---|---|---|---|---|---|---|
| Control task | (9a) | (S16a) | (S16b) | (S16c) | (9b) | (9c) | (S16d) | (9d) |
| $j$ | 1 | | | | | | | |
| $\mathcal{H}$ | $[10^{-5}, 5]^4$ | | | | | | | |
| $h_1$ | 1 | 2 | 3 | 4 | 1 | 3 | 3 | 3 |
| $h_2$ | 1 | 2 | 3 | 4 | 2 | 2 | 2 | 1 |
| $h_3$ | 1 | 2 | 3 | 4 | 2 | 1 | 1 | 3 |
| $h_4$ | 1 | 2 | 3 | 4 | 3 | 0.5 | 3 | 1 |

Table S14: Parameters used to define the optimisation program for the controlled Production and Degradation Chemical Reaction Network.

| Controlled Production and Degradation | | | | | | | | |
|---|---|---|---|---|---|---|---|---|
| Control task | (9a) | (S16a) | (S16b) | (S16c) | (9b) | (9c) | (S16d) | (9d) |
| $n_{\text{iter}}$ | 20000 | 20000 | 20000 | 20000 | 20000 | 30000 | 20000 | 20000 |
| $\gamma$ | 0.01 | | | | | | | |
| $\varepsilon$ | $10^{-7}$ | $10^{-7}$ | $10^{-7}$ | $10^{-7}$ | $10^{-7}$ | $10^{-8}$ | $10^{-7}$ | $10^{-7}$ |
| $N_{\max}$ | 200 | | | | | | | |

Table S15: Hyperparameters of the Projected Gradient Descent for the controlled Production and Degradation Chemical Reaction Network.

| Controlled Bursting Gene | | | | | | | | |
|---|---|---|---|---|---|---|---|---|
| Control task | (S17a) | (10a) | (S17b) | (S17c) | (10b) | (S17d) | (10c) | (10d) |
| $j$ | 2 | | | | | | | |
| $\mathcal{H}$ | $[10^{-10}, 5]^4$ | | | | | | | |
| $h_1$ | 0.5 | 1 | 1.5 | 1.75 | 0.5 | 0.5 | 1.75 | 1.5 |
| $h_2$ | 0.5 | 1 | 1.5 | 1.75 | 1 | 1 | 1.5 | 0.5 |
| $h_3$ | 0.5 | 1 | 1.5 | 1.75 | 1 | 1.5 | 1 | 1.5 |
| $h_4$ | 0.5 | 1 | 1.5 | 1.75 | 1.5 | 1.75 | 0.5 | 0.5 |

Table S16: Parameters used to define the optimisation program for the controlled Bursting Gene Chemical Reaction Network.

| Controlled Bursting Gene | | | | | | | | |
|---|---|---|---|---|---|---|---|---|
| Control task | (S17a) | (10a) | (S17b) | (S17c) | (10b) | (S17d) | (10c) | (10d) |
| $n_{\text{iter}}$ | 30000 | 30000 | 30000 | 50000 | 30000 | 30000 | 30000 | 30000 |
| $\gamma$ | 0.01 | | | | | | | |
| $\varepsilon$ | $10^{-8}$ | | | | | | | |
| $N_{\max}$ | 200 | | | | | | | |

Table S17: Hyperparameters of the Projected Gradient Descent for the controlled the Bursting Gene Chemical Reaction Network.

| Toggle Switch | | | | | | | | |
|---|---|---|---|---|---|---|---|---|
| Control task | (S18a) | (S18b) | (11a) | (S18c) | (11b) | (S18d) | (11c) | (11d) |
| $j$ | 2 | | | | | | | |
| $\mathcal{H}$ | $[10^{-5}, 5]^4$ | | | $[10^{-5}, 4]^4$ | | | | |
| $h_1$ | 1.5 | 2 | 3 | 4 | 2 | 1.5 | 4 | 4 |
| $h_2$ | 1.5 | 2 | 3 | 4 | 3 | 2 | 3 | 1 |
| $h_3$ | 1.5 | 2 | 3 | 4 | 3 | 3 | 2 | 4 |
| $h_4$ | 1.5 | 2 | 3 | 4 | 4 | 4 | 1 | 1 |

Table S18: Parameters used to define the optimisation program for the Toggle Switch Chemical Reaction Network.

| Toggle Switch | | | | | | | | |
|---|---|---|---|---|---|---|---|---|
| Control task | (S18a) | (S18b) | (11a) | (S18c) | (11b) | (S18d) | (11c) | (11d) |
| $n_{\text{iter}}$ | 50000 | 30000 | 30000 | 30000 | 50000 | 20000 | 50000 | 50000 |
| $\gamma$ | 0.001 | 0.005 | 0.001 | 0.001 | 0.005 | 0.01 | 0.005 | 0.005 |
| $\varepsilon$ | $10^{-7}$ | $10^{-9}$ | $10^{-9}$ | $10^{-8}$ | $10^{-7}$ | $10^{-7}$ | $10^{-7}$ | $10^{-7}$ |
| $N_{\max}$ | 200 | | | | | | | |

Table S19: Hyperparameters of Projected Gradient Descent for the Toggle Switch Chemical Reaction Network.

Note that $N_{\max}$ could be made to vary with the value of the parameters at the current descent iteration.
